# *Candida glabrata* replicating within macrophages experiences amino acid deprivation, DNA damage, and chromosome instability

**DOI:** 10.1101/2025.09.15.676333

**Authors:** Ariel A. Aptekmann, Nathaly Cabrera, Melody C. Hayman, Ruth Watson, Joseph A. Stewart, Christopher Quinteros, Chang Liu, Caroline E. Hayter, Mikhail V. Keniya, David S. Perlin, Juan Lucas Argueso, Erika Shor

## Abstract

Macrophages, the central players of innate immunity, control invading microbes by encapsulating them inside the phagosome, a nutrient-poor, reactive oxidant species-rich organelle. Nevertheless, some microbes, including the opportunistic yeast pathogen *Candida glabrata*, noted for its karyotype diversity, rapid evolution of antifungal drug resistance, and lack of meiosis, can survive and even replicate inside macrophages. However, it is not fully understood how *C. glabrata* responds to macrophage engulfment, and it is unknown how this presumably DNA-damaging environment influences the pathogen’s genome stability. In this study, we used comparative transcriptomics to identify amino acid starvation and DNA damage as conditions eliciting *C. glabrata* responses most similar to macrophage engulfment. Consistent with this, we found that *C. glabrata* intra-macrophage survival and replication require master regulator of amino acid biosynthesis *GCN4* and functional DNA double-strand break repair. Furthermore, comet assays provided the first direct evidence for increased DNA breaks in intra-macrophage yeast, and pulse-field gel electrophoresis showed that chromosomal alterations occur frequently in macrophage-passaged *C. glabrata*. Interestingly, these alterations could not be resolved by long read DNA sequencing, suggesting that they involved highly complex repetitive regions. Finally, we identified several point mutations emerging during macrophage passaging and showed that among them, a frameshift in *RME1* (repressor of <u>me</u>iosis in *Saccharomyces cerevisiae*), increased *C. glabrata* intra-macrophage fitness. Together, these analyses point to amino acid deprivation, reveal elevated DNA breakage and chromosome instability, and raise intriguing questions about the role of meiotic gene orthologs in *C. glabrata* persisting and replicating within macrophages.

## Introduction

Macrophages are critical components of innate immunity, serving as the first line of defense against pathogens. Upon encountering a microbe, macrophages envelop it inside a specialized organelle, the phagosome, and attempt to destroy it using multiple means, including low pH, reactive oxidant (oxygen and nitrogen) species (ROS), and severe nutrient limitation. Thus, a key feature of a successful pathogen is the ability to avoid macrophage-mediated killing. Fungal pathogens, which cause over 1.7 million deaths per year globally (1), possess multiple mechanisms to withstand and escape macrophages (2, 3). Upon being encapsulated in the phagosome, fungi upregulate nutrient acquisition and biosynthesis pathways, often switching from glycolytic to gluconeogenic metabolism and using alternative carbon sources, such as amino acids or fatty acids (4–8). Some fungi taken up by macrophages induce filamentation, rupture macrophages from the inside, and escape (2, 3). Finally, to protect themselves from macrophage-generated ROS, fungi upregulate ROS-detoxifying enzymes, such as catalase, and interfere with phagosome maturation, resulting in reduced acidification and ROS production (3). Although ROS are a key feature of the intra-macrophage environment, their effects on the engulfed pathogens are relatively poorly understood. In particular, although it has been routinely postulated that the ROS damage the microbe’s DNA, direct evidence for this is lacking, and no studies have examined the consequences of macrophage engulfment on the pathogen’s genome stability.

*Candida glabrata* (taxonomic name *Nakaseomyces glabratus*), the second most frequent cause of invasive candidiasis after *C. albicans*, presents a clinical challenge due to its rapid development of antifungal drug resistance (9, 10). *C. glabrata* has also been noted for its remarkable ability to survive and even replicate within macrophages, suggesting that they constitute an important host niche for this pathogen (11, 12). Unlike *C. albicans*, *C. glabrata* is a non-dimorphic yeast that does not form filaments but persists inside macrophages in yeast form. Several studies have examined gene expression (6, 13) or RNA polymerase gene occupancy (7) in *C. glabrata* taken up by macrophages. As in other fungi, results have revealed extensive metabolic reprogramming to upregulate nutrient acquisition and reroute carbon metabolism, as well as upregulation of ROS detoxifying genes (6, 7, 13). Interestingly, some DNA damage checkpoint genes were also upregulated in intra-macrophage *C. glabrata* (7), consistent with a DNA-damaging environment. Also, transposon screens and targeted mutant analysis found that *C. glabrata* intra-macrophage fitness decreases upon deletion of several genes involved in DNA damage repair (13). However, the genes involved, such as *CgSGS1*, *CgRTT107*, and *CgRTT109*, affect multiple DNA damage repair pathways, leaving it unclear what types of DNA damage might be at play. Thus far, no direct evidence of damage (e.g., DNA breaks) has been reported in intra-macrophage *C. glabrata* or, to the best of our knowledge, in any other macrophage-engulfed pathogen.

In several fungal pathogens, the host environment appears to trigger genetic instability. This phenomenon is best-studied in *C. albicans*, where strains isolated from patients or passed through animals often carry whole chromosome or segmental aneuploidies, copy number variants, and other genome rearrangements (14–16). *C. glabrata* is more closely related to the nonpathogenic baker’s yeast *Saccharomyces cerevisiae* than to *C. albicans* and is predominantly haploid with few reports of aneuploidy. Interestingly, although *C. glabrata* genome contains orthologs of *S. cerevisiae* mating and meiosis genes, it does not appear to mate or sporulate, at least under laboratory conditions. However, a hallmark feature of *C. glabrata* is the great diversity of karyotypes, or chromosomal configurations, observed in different clinical strains (17–19). Because there is no *C. glabrata* analog of the *S. cerevisiae* gross chromosome rearrangement assay (20), the causes of chromosomal instability in this yeast have not been systematically examined. In *S. cerevisiae*, some of the greatest rates of chromosome instability are seen in mutants lacking functional DNA damage or DNA replication checkpoints (20–22). Interestingly, we have reported that compared to *S. cerevisiae*, *C. glabrata* has an attenuated DNA damage checkpoint (23), suggesting that a DNA damaging environment, such as within the macrophage, may trigger genome instability in this organism. However, this hypothesis has not yet been tested.

To study the stress responses, DNA integrity, and genome stability of macrophage-engulfed *C. glabrata*, we used comparative transcriptomics, deletion mutant analysis, comet assays, pulse-field gel electrophoresis, and whole genome sequencing. We discovered transcriptional signatures of amino acid deprivation, presence of increased DNA breaks, and frequent chromosome rearrangements in *C. glabrata* persisting and replicating within macrophages. Interestingly, we also identified an instance of evolution of higher intra-macrophage fitness associated with a loss of function mutation in a negative regulator of meiosis, raising the possibility that the meiotic program may play a role in intra-macrophage *C. glabrata*. Together, these results shed new light on how host innate immunity shapes genome configuration and evolution of this important opportunistic fungal pathogen.

## Results

### Selection of stress conditions for cultured *C. glabrata*

We reasoned that comparing the transcriptional response of *C. glabrata* to macrophage engulfment to its responses to different *in vitro* stresses may shed new light on the intra-macrophage environment. Thus, for our transcriptomic analysis, we selected a diverse panel of stresses: oxidative (H_2_O_2_), DNA damage (MMS), osmotic (NaCl), cell wall (Congo Red), acidic pH, carbon starvation, nitrogen starvation, and amino acid starvation. First, growth curves were performed to identify the concentrations of H_2_O_2_, MMS, NaCl, Congo Red, and pH that result in roughly similar effects on *C. glabrata* growth (Fig. 1A). This was not done for carbon or nitrogen starvation, which did not support growth, or for amino acid starvation because the strain (CBS138) is prototrophic. We observed that different stresses affected *C. glabrata* growth with different dynamics: e.g., cells shifted to acidic pH grew well at first but then slowed down, whereas cells exposed to Congo Red grew slowly at first but after several hours adapted and grew better (Fig. 1A). The concentrations selected for RNA sequencing (RNAseq) and their effects on *C. glabrata* growth are shown in Figure 1A. For most stress conditions, cells were shifted into the stressor medium and collected at 1h, 2h, and 4h post-shift. For the amino acid starvation condition, cells were collected at 1h and 4h post-shift. Finally, control samples were collected at 0h (pre-stress) and from non-stressed cultures at 1h, 2h, and 4h to ensure adequate normalization of the resulting RNA signals.

**Figure 1.**
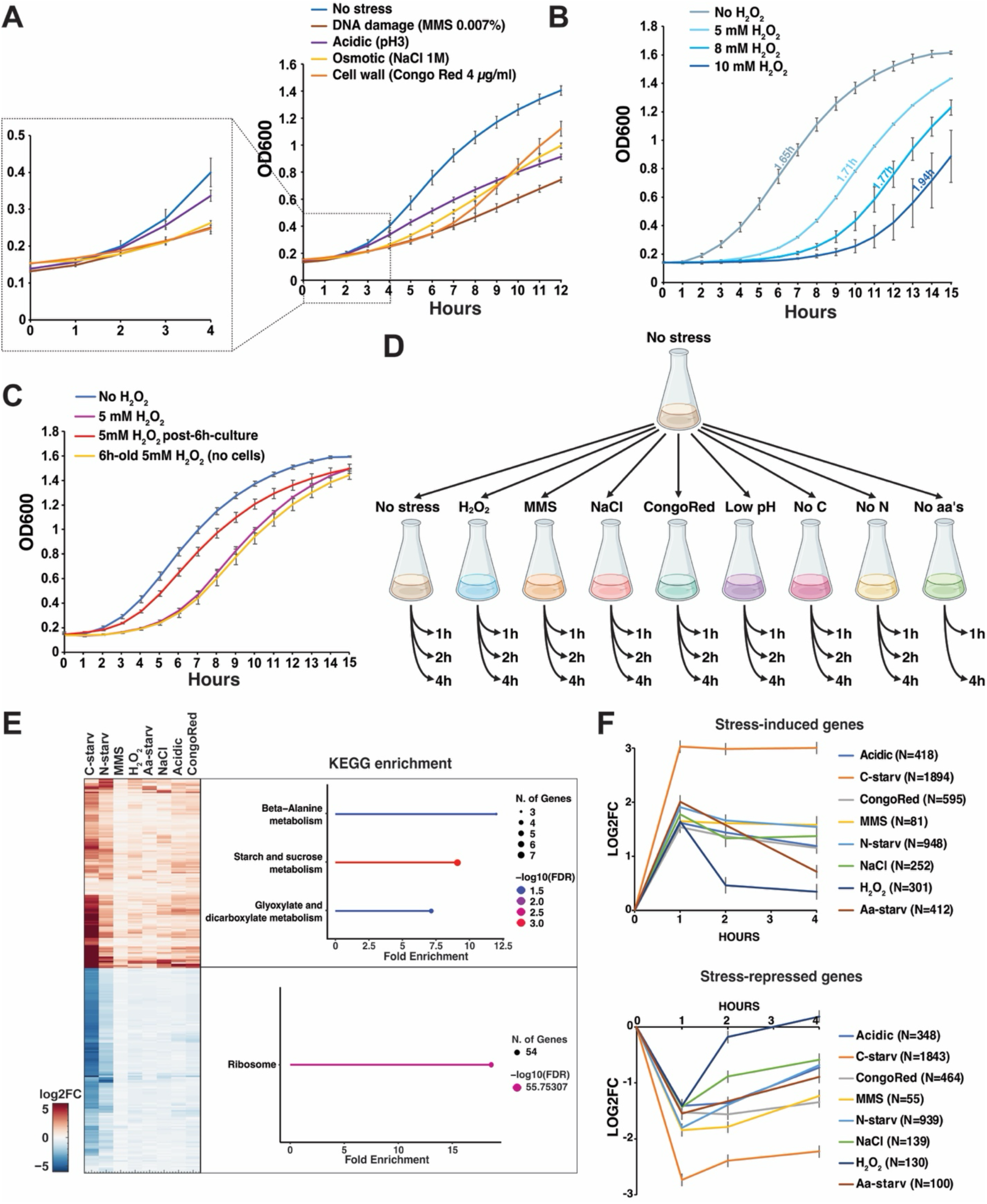
*C. glabrata* transcriptional responses to a diverse panel of stresses reveal a core set of stress-induced and stress-repressed genes and different stress adaptation dynamics. **A.** Growth curves of *C. glabrata* strain CBS138 in SD medium containing the indicated stressors. Cells for RNAseq were harvested within 4 hours of stress exposure. **B.** Effects of increasing H_2_O_2_ concentrations in *C. glabrata* growth. The numbers indicate doubling times once cells resumed growth. **C.** 5 mM H_2_O_2_ causes a lag in *C. glabrata* growth (purple), but if the H_2_O_2_ is first detoxified by *C. glabrata* (red), no lag is observed. **D.** Sample collection scheme for RNAseq. This panel was created with the help of Biorender (institutional license). **E.** KEGG enrichment categories for genes induced or repressed by all examined stresses. The heatmap was generated using InstantClue v0.12.2 (64) and the enrichment plot was generated using ShinyGO V0.81 (65). F. *C. glabrata* cells adapt to H_2_O_2_ stress much more rapidly than to other stresses. Growth curve results represent an average of three or more independent biological replicates. Error bars represent the standard deviation.

Exposure to H_2_O_2_ produced an unique effect where the addition of increasing concentrations of H_2_O_2_ increased lag times prior to growth resumption much more strongly than the growth rate (doubling time) once growth resumed (Fig. 1B). We hypothesized that this was because *C. glabrata* resumed growth only after nearly completely detoxifying the H_2_O_2_. To test this hypothesis, we cultured *C. glabrata* in synthetic medium containing 5mM H_2_O_2_ for 6 hours, then filtered out the cells and added new cells to the medium. Consistent with the prediction that the first batch of cells had detoxified the H_2_O_2_, growth of the newly added cells was only mildly slowed compared to fresh medium, and this mild slowing was likely due to the fact that the first batch of cells had somewhat depleted the nutrients (Fig. 1C). In contrast, fresh medium containing 5mM H_2_O_2_ as well as 6h old medium containing 5mM H_2_O_2_ (to control for potential H_2_O_2_ degradation) produced the stereotypical lag times (Fig. 1C). Based on these experiments, to ensure that *C. glabrata* experiences strong and consistent oxidative stress during the 4h experimental time-course, we exposed *C. glabrata* to 10mM H_2_O_2_, collected cells at 1h and 2h, spiked the cultures with additional 10mM H_2_O_2_ at 2h, and collected the last set of samples at 4h.

### Transcriptional responses of *C. glabrata* to a panel of different environmental stressors reveal a common core stress response and differing adaptation dynamics

Total RNA was isolated from all samples (Fig. 1D) and sequenced at Azenta Life Sciences. Our differential expression analysis generated Log2 fold-change (LFC) values and corresponding adjusted p-values to assess the significance of each treatment’s effect at different time points relative to non-stressed controls at baseline (Dataset S1). Our experimental strategy of exposing *C. glabrata* to a diverse panel of stresses and of collecting samples over 4 hours post-shift to stress condition has allowed us to ask two questions: first, whether we can detect a core acute transcriptional stress response common to all stresses, and second, whether transcriptional adaptation to stress proceeds differently for different stresses over the 4-hour timeframe.

To address the first question, we identified sets of genes that were significantly (FDR p<0.05) induced or repressed under all stress conditions at 1 hour post-stress exposure (Fig. 1E). We identified 132 genes significantly upregulated by all stresses, of which all but five (CAGL0B02981g, CAGL0B03014g, CAGL0H00759g, CAGL0L00205g, and CAGL0E03366g) had orthologs in *S. cerevisiae*, and 143 genes significantly downregulated by all stresses, of which all but one (CAGL0M02321g) had orthologs in *S. cerevisiae* (Table S1). Using String-DB and KEGG enrichment analyses, we found that both upregulated and downregulated gene sets formed highly interconnected networks (Fig. S1, Fig. S2) enriched for certain pathways (Fig. 1E). Stress-upregulated genes were highly enriched for genes functioning in carbon metabolism, specifically starch and sucrose metabolism, beta-alanine metabolism, and glyoxylate and dicarboxylate metabolism (Fig. 1E). Additionally, stress-upregulated genes contained several functional clusters that did not meet the KEGG significance threshold but may still be informative, such as clusters of genes involved in fatty acid catabolism, autophagy, iron-sulfur cluster maturation, pheromone signaling, and non-homologous end joining (Fig. S1). These results are consistent with *C. glabrata* turning up catabolic pathways and the use of alternative and internal energy sources upon encountering environmental stress. On the other hand, the universal upregulation of the α-factor gene MFα and of *STE3*, which encodes the a-factor receptor (CBS138 belongs to the α mating type), in response to stress is intriguing because *C. glabrata* has never been observed to mate. The stress-downregulated gene set was strongly enriched for ribosome components and other genes functioning in cytoplasmic translation (Fig. 1E, Fig. S2), which is very similar to results obtained in *S. cerevisiae* (24) and indicates that downregulation of translation is a common response of *C. glabrata* to diverse stresses.

To address the second question (i.e., the dynamics of transcriptional adaptation to stress), we focused on sets of genes strongly (log2FC>1, FDR p<0.05) induced or repressed by each stress at 1 hour post-stress exposure and asked how their expression changed over the 4-hour time course. This analysis produced several insights. First, the transcriptional response to carbon starvation was the strongest and most sustained, involving the largest number of genes, the greatest levels of induction or repression, and little transcriptional adaptation, i.e., dampening of the transcriptional changes by 4 hours, particularly for carbon starvation-induced genes (Fig. 1F). For most other stresses (nitrogen starvation, osmotic stress, DNA damage, cell wall stress, and acidic stress), there was a slight dampening of gene induction, and a moderate to strong dampening of gene repression by 4 hours (Fig. 1F), consistent with adaptation to stress. For amino acid starvation, the transcriptional response was strongly attenuated by 4 hours, consistent with the prototrophic CBS138 strain adapting well to amino acid starvation, likely by turning on amino biosynthesis pathways. Interestingly, the strongest attenuation of the transcriptional response, both for induced and repressed genes, was observed for oxidative stress (Fig. 1F), despite the high dose of H_2_O_2_ (10 mM) and the spiking of the culture with additional H_2_O_2_ at 2 hours. This observation underscores the previously noted exquisite adaptation of *C. glabrata* to highly oxidative environments, such as host macrophages (12, 25).

### *C. glabrata* transcriptional response to the intra-macrophage environment most closely resembles its *in vitro* transcriptional responses to amino acid starvation and DNA damage

We previously generated RNAseq data from *C. glabrata*-infected THP-1 cells to study how mitochondrial disfunction (petiteness) (26) or drug resistance (27) of *C. glabrata* affect interactions with macrophages. Thus, we now had the opportunity to compare the transcriptional response of wild type *C. glabrata* (reference strain CBS138) to macrophage engulfment, for which we had data from two time-points (3 and 24 hours post-infection) to those induced by a variety of *in vitro* stress conditions up to 4 hours after stress exposure. We compared the transcriptomic datasets using two methods: Principal Component Analysis (Fig. 2A) and Pearson correlation coefficient (Fig. 2B). For the latter, we built a heatmap based on the sample-to-sample correlation coefficient and agglomerative clustering was used to sort the samples and generate a heatmap and a dendrogram. The two methods produced several convergent insights. First, both methods indicated that acutely stressed *C. glabrata* in culture is more similar in terms of transcription to intra-macrophage *C. glabrata* at 3 hours post-infection than at 24 hours post-infection, consistent with the conclusion that *C. glabrata* experiences certain environmental stresses upon macrophage engulfment and then adapts to those stresses. Second, we observed that the transcriptional changes characteristic of the 3-hour post-macrophage engulfment timepoint were most similar to those elicited by amino acid starvation and DNA damage (Fig. 2A, 2B). This conclusion was reinforced by additional analysis of the genes strongly (log2FC>1, FDR p<0.05) induced in *C. glabrata* at 3 hours post-macrophage infection. KEGG pathway analysis identified multiple amino acid biosynthesis pathways as being significantly enriched in the macrophage-induced gene set (Fig. 2C). Likewise, String-DB analysis identified several highly interconnected clusters in this gene set, the largest of which was comprised of amino acid biosynthesis genes, whereas the second largest cluster contained genes whose orthologs in *S. cerevisiae* function in meiotic DNA double-strand break (DSB) formation and meiotic recombination (*SPO11*, *SPO13*, *SPO22*, *ZIP1*, *GMC2*, *MEI1*, *MLH3*, *SAE3*, and *REC8*), resolution of recombination intermediates (*MUS81*, *NSE3*, *NSE5*, and *KRE29*), checkpoint activation (*CHK1*, *DDC1*, *PCH2*), and telomere maintenance (*EST2*, *RIF2*, *RRM3*, *SAE2* and *TEL2*) (Fig. S3). Whereas *C. glabrata* has not been observed to undergo meiosis, the upregulation of these genes was consistent with activation of DNA damage responses and the observed similarity between *C. glabrata* transcriptional responses to macrophages and to MMS (Fig. 2A).

**Figure 2.**
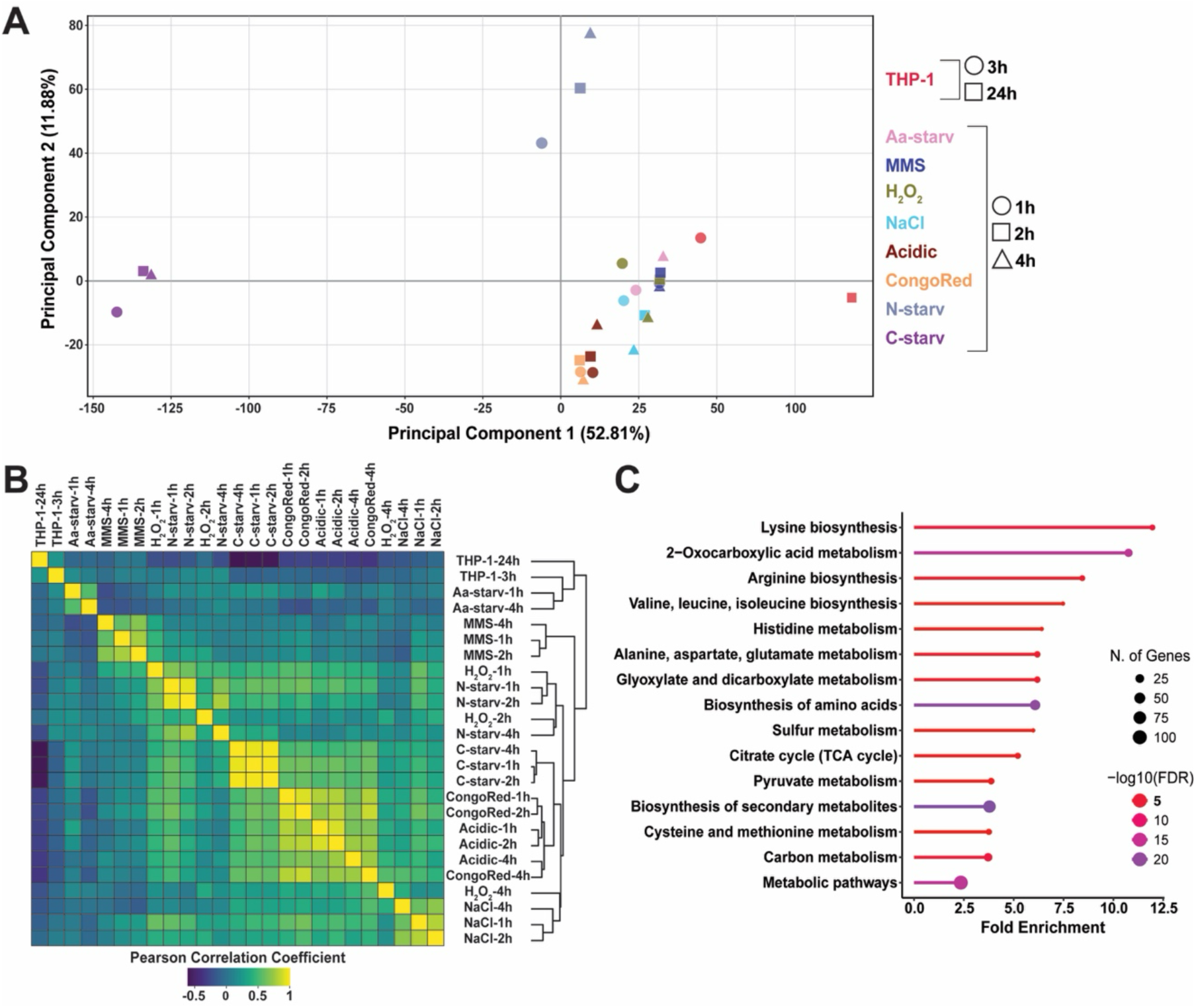
*C. glabrata* transcriptional response to macrophage engulfment most closely resembles its responses to amino acid deprivation and DNA damage. **A.** Principal Component Analysis of the transcriptional responses to the diverse *in vitro* stresses and to macrophages. **B.** Pairwise Pearson correlation coefficients among the transcriptional responses to the diverse *in vitro* stresses and to macrophages. **C.** KEGG enrichment categories for *C. glabrata* genes induced (>2-fold, FDR<0.05) at 3 hours after macrophage infection. The plot was generated using ShinyGO (65).

### *C. glabrata* intra-macrophage fitness requires *GCN4*, the master transcriptional regulator of amino acid metabolism

The upregulation of amino acid biosynthesis genes in intra-macrophage *C. glabrata* and its similarity to the transcriptional response to amino acid starvation suggested that the phagosome constituted an amino acid-poor environment. Thus, we hypothesized that *C. glabrata* unable to induce amino acid biosynthesis genes in response to amino acid scarcity would lose intra-macrophage fitness. To test this hypothesis, we deleted *GCN4*, which encodes a master transcriptional regulator of amino acid biosynthesis (28). Consistent with the role of its orthologs in other organisms, deletion of *GCN4* resulted in the inability of *C. glabrata* to grow in the absence of exogenous amino acids but did not affect the yeast’s sensitivity to oxidative stress or DNA damage (Fig. 3A). We also asked whether *GCN4* is required for *C. glabrata* survival in the absence of carbon or nitrogen by seeding cells/ml in synthetic medium lacking either glucose or ammonium sulfate, culturing for 24 hours, and plating for CFU. Whereas we found no effect of the *gcn4Δ* mutant in the absence of nitrogen, *gcn4Δ* slightly but significantly reduced *C. glabrata* survival in the absence of a carbon source (Fig. 3B). Finally, we infected THP-1 macrophages with the WT control and the *gcn4Δ* mutant. Consistent with the phagosome being an amino acid-poor environment, we found that deletion of *GCN4* significantly reduced *C. glabrata* intra-macrophage fitness, measured as the increase in intra-macrophage CFU over 24 hours of infection (Fig. 3C). These results supported the conclusion that *C. glabrata* needs to up-regulate amino acid biosynthesis to survive inside macrophages, suggesting that it is an amino acid depleted environment.

**Figure 3.**
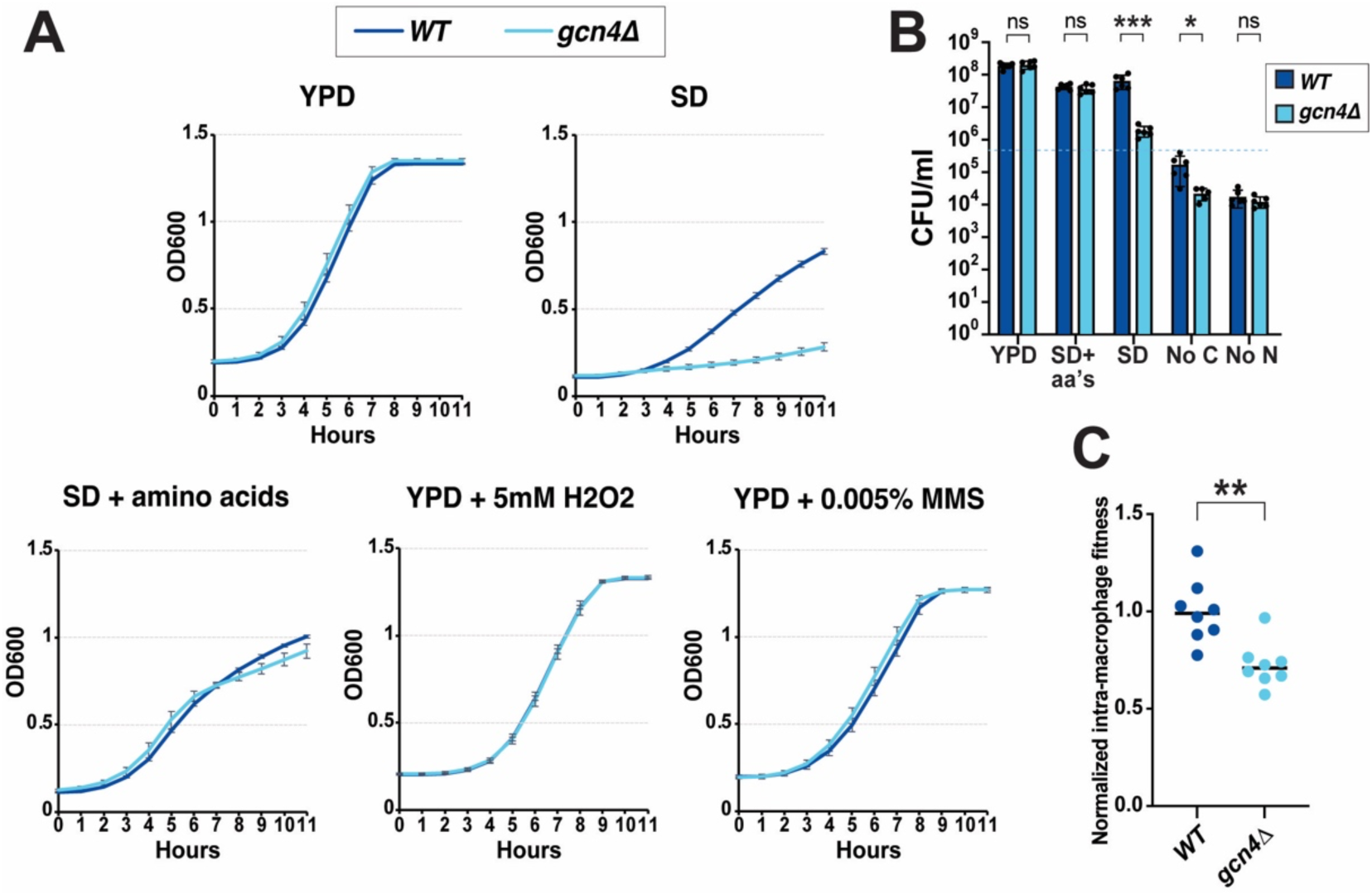
Deletion of master regulator of amino acid biosynthesis *GCN4* reduces *C. glabrata* intra-macrophage fitness. **A.** The *gcn4Δ* mutant is impaired for growth in the absence of exogenous amino acids but not in rich medium, in synthetic medium supplemented with amino acids, or in the presence of oxidative stress or DNA damage. Growth curve results represent an average of three or more independent biological replicates. Error bars represent the standard deviation. **B.** Wild type and *gcn4Δ* cells were inoculated into the indicated media at 5×10^5^ cells/ml (indicated by the dashed line), cultured for 24 hours, and the resulting CFU were plotted. **C.** The *gcn4Δ* mutant had reduced intra-macrophage fitness, calculated as the fold-increase in intra-macrophage yeast CFU between 3h and 24h post-infection normalized to the wild-type control. All data were normally distributed and t-tests were used for pairwise comparisons. Each dot represents a biological replicate. * p<0.05, ** p<0.01, *** p<0.001.

### Intra-macrophage *C. glabrata* experiences DNA breaks and relies on DNA double-strand break repair mechanisms

It has long been postulated that the phagosome is a DNA-damaging environment due to the release of ROS; however, direct evidence for elevated DNA damage in the engulfed microbes has been lacking. To directly assess the levels of DNA breaks (both single and double strand) in macrophage-engulfed *C. glabrata*, we used alkaline single-cell gel electrophoresis, a.k.a. the comet assay (29). The comet assay is widely used in mammalian cells to measure DNA breaks, but has been more rarely used in microbes, including yeast, possibly because microbial cells contain much less DNA and the signal from single cells is much weaker. We optimized the comet assay for *C. glabrata* (see Methods) and were able to detect single cell nuclear DNA fluorescence upon staining of agarose-embedded lysed cells with propidium iodide (PI) (Fig. 4A). We used the OpenComet tool (30) to characterize comets produced by *C. glabrata* cells obtained from THP-1 macrophages 24 hours after infection, as well as control *C. glabrata* cultured in the same medium as macrophages (RPMI) for 24 hours. Cells cultured in RPMI produced predominantly spherical DNA signals with no tails, indicating largely undamaged DNA (Fig. 4B). On the other hand, OpenComet analysis identified an increased number of comets with small tails in cells obtained from macrophages, consistent with increased levels of DNA breaks in *C. glabrata* residing within macrophages (Fig. 4B). Because dying and dead cells have fragmented DNA, we asked whether *C. glabrata* cells within macrophages might be in the process of cell death, thus accounting for the enhanced comet signal. However, there was no significant difference between the plating efficiencies of RPMI-cultured and intra-macrophage *C. glabrata* (Fig. 4C), indicating that the increased comet signal was due to increased DNA breaks in living cells.

**Figure 4.**
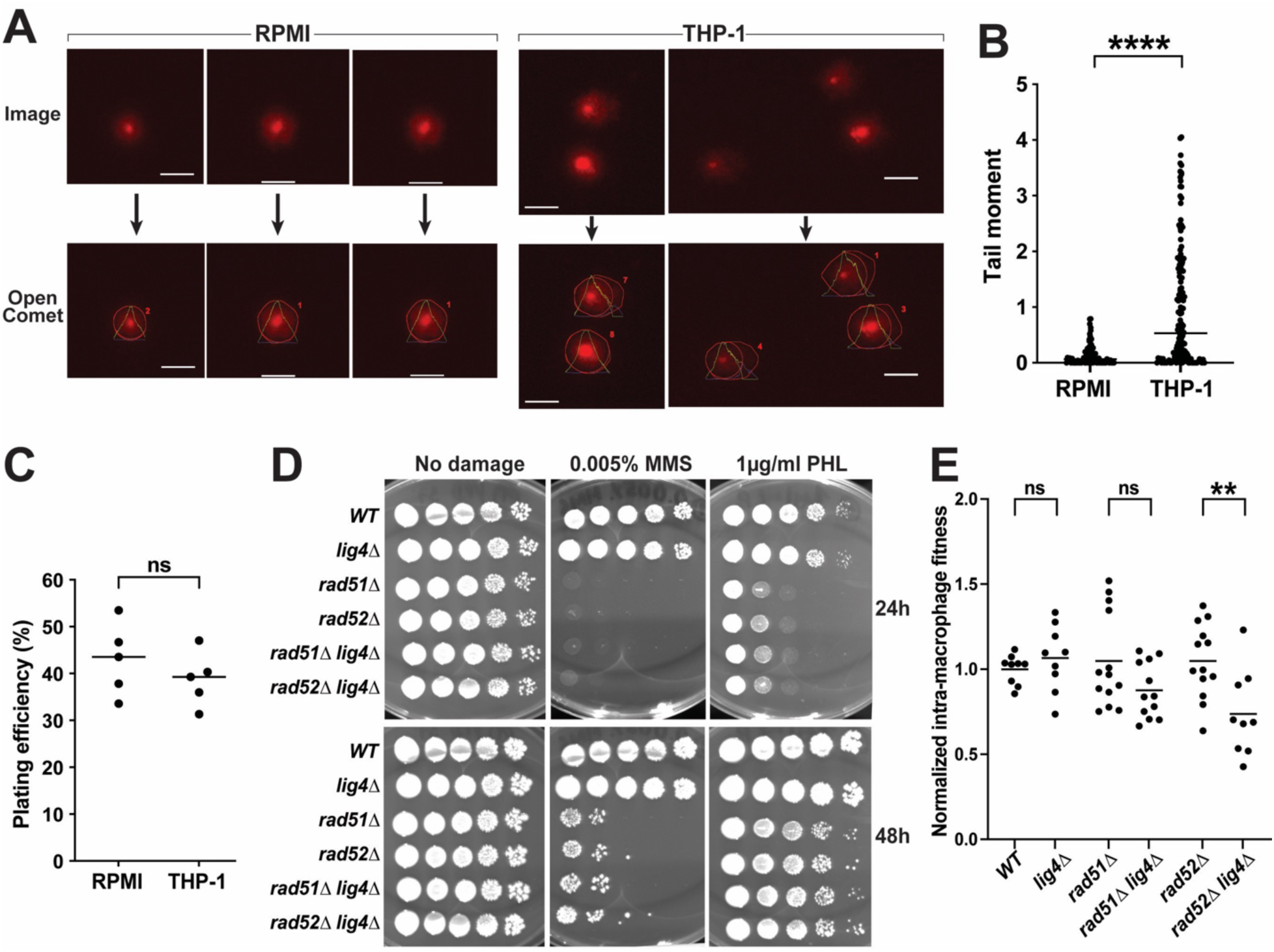
Intra-macrophage *C. glabrata* has elevated levels of DNA breaks and requires DNA double-strand break repair for optimal intra-macrophage fitness. **A.** Examples of images of comets and their analysis by OpenComet (30). Bar = 10 µm. **B.** Tail moment analysis identified more DNA breaks in *C. glabrata* from macrophages than *C. glabrata* cultured in RPMI. The tail moment was calculated by OpenComet by multiplying the percentage of DNA in the comet tail by the length of the comet tail (30). **C.** RPMI-cultured and intra-macrophage *C. glabrata* have similar plating efficiency, indicating similar viability. Plating efficiency was calculated by counting cells under a microscope, plating a defined cell number, and dividing the resulting CFU by the number of plated cells. **D.** HR mutants *rad51Δ* and *rad52Δ* are sensitive to DNA damaging agents methyl methanesulfonate (MMS) and phleomycin (PHL), whereas deletion of NHEJ gene *LIG4* had no effect. **E.** Deletion of both HR gene *RAD52* and NHEJ gene *LIG4* synergistically reduces *C. glabrata* intra-macrophage fitness, calculated as the fold-increase in intra-macrophage *C. glabrata* between 3h and 24h post-infection normalized to the wild-type control. Each dot represents a biological replicate. For pairwise comparisons, the t-test was used when data was normally distributed and the Mann-Whitney test was used when data was not normally distributed. * p<0.05, ** p<0.01, *** p<0.001, **** p<0.0001.

Our transcriptional data suggested that upon macrophage engulfment *C. glabrata* upregulates genes whose *S. cerevisiae* orthologs function in the formation and processing of meiotic DSBs (Fig. S3). DSBs are considered some of the most toxic DNA lesions, which can lead to genome rearrangements and cell death (31). Cells can repair DSBs using two alternative mechanisms: homologous recombination (HR), which relies on the presence of a homologous template (e.g., sister chromatid or homologous chromosome) and is relatively error-free, and non-homologous end joining (NHEJ), which ligates broken ends together without using a template and is considered more error-prone (31). To address the importance of DSB repair in intra-macrophage *C. glabrata*, we created deletions of HR genes *RAD51* and *RAD52* and NHEJ gene *LIG4* (a.k.a. *DNL4*). We found that *in vitro*, *rad51Δ* and *rad52Δ* strains, but not *lig4Δ*, were sensitive to MMS and DNA break-inducing agent phleomycin, consistent with their phenotypes in *S. cerevisiae*. Interestingly, impairing either HR or NHEJ separately did not reduce the ability of *C. glabrata* to proliferate within macrophages (Fig. 4E). However, impairing both pathways by deleting *RAD52* and *LIG4* resulted in a synergistic reduction in intra-macrophage fitness (Fig. 4E). A weaker, non-statistically significant reduction was observed in the *rad51Δ lig4Δ* mutant. Interestingly, no synergy was observed upon DNA damage *in vitro*, as deleting both *RAD51* and *LIG4* or both *RAD52* and *LIG4* did not result in additional sensitivity to MMS or PHL (Fig. 4D). We conclude that intra-macrophage *C. glabrata* forms DSBs that can be repaired either by HR or NHEJ, whereas the DSBs caused in cultured cells by MMS or phleomycin are repaired predominantly by HR.

### Intra-macrophage *C. glabrata* experiences chromosomal instability

Formation of DNA DSBs triggers the activation of checkpoint signaling pathways that facilitate DSB repair and prevent genome instability (32). In *S. cerevisiae*, deletion of the central DNA damage checkpoint kinase *RAD53* and other checkpoint components leads to increased gross chromosomal rearrangements (20–22). We previously reported that *C. glabrata* Rad53 activity is attenuated relative to its *S. cerevisiae* ortholog (23), suggesting that *C. glabrata* may be more prone to chromosomal rearrangements upon DNA damage – for instance, within the macrophage phagosome. To test this hypothesis, we serially passaged *C. glabrata* cells through macrophages three times and analyzed the karyotypes of 15 randomly chosen colonies from every passage and also from the starting culture (Fig. 5A). After every passage, macrophages were lysed and the yeast collected and immediately used to re-infect a fresh batch of macrophages without any outgrowth in culture medium. Furthermore, no bottleneck events were introduced, as the entirety of the yeast collected from macrophages were used to reinfect the next batch (with the exception of a small fraction of cells plated on YPD for CFU enumeration and collection of 15 random colonies). After each 48-hour passage, the number of *C. glabrata* increased, consistent with intra-macrophage proliferation, and the final number of cells obtained after three passages was ∼25-fold higher than the number used in the first infection (Fig. 5B). In total, we collected 60 isolates: 15 from the original culture (RPMI-1 through RPMI-15), 15 from the first passage (P1-1 through P1-15), 15 from the second passage (P2-1 through P2-15), and 15 from the third passage (P3-1 through P3-15). Thus, isolates from RPMI had never experienced the intra-macrophage environment, whereas, at the other end, P3 isolates had been successively taken up by, and proliferated within, three sets of macrophages.

**Figure 5.**
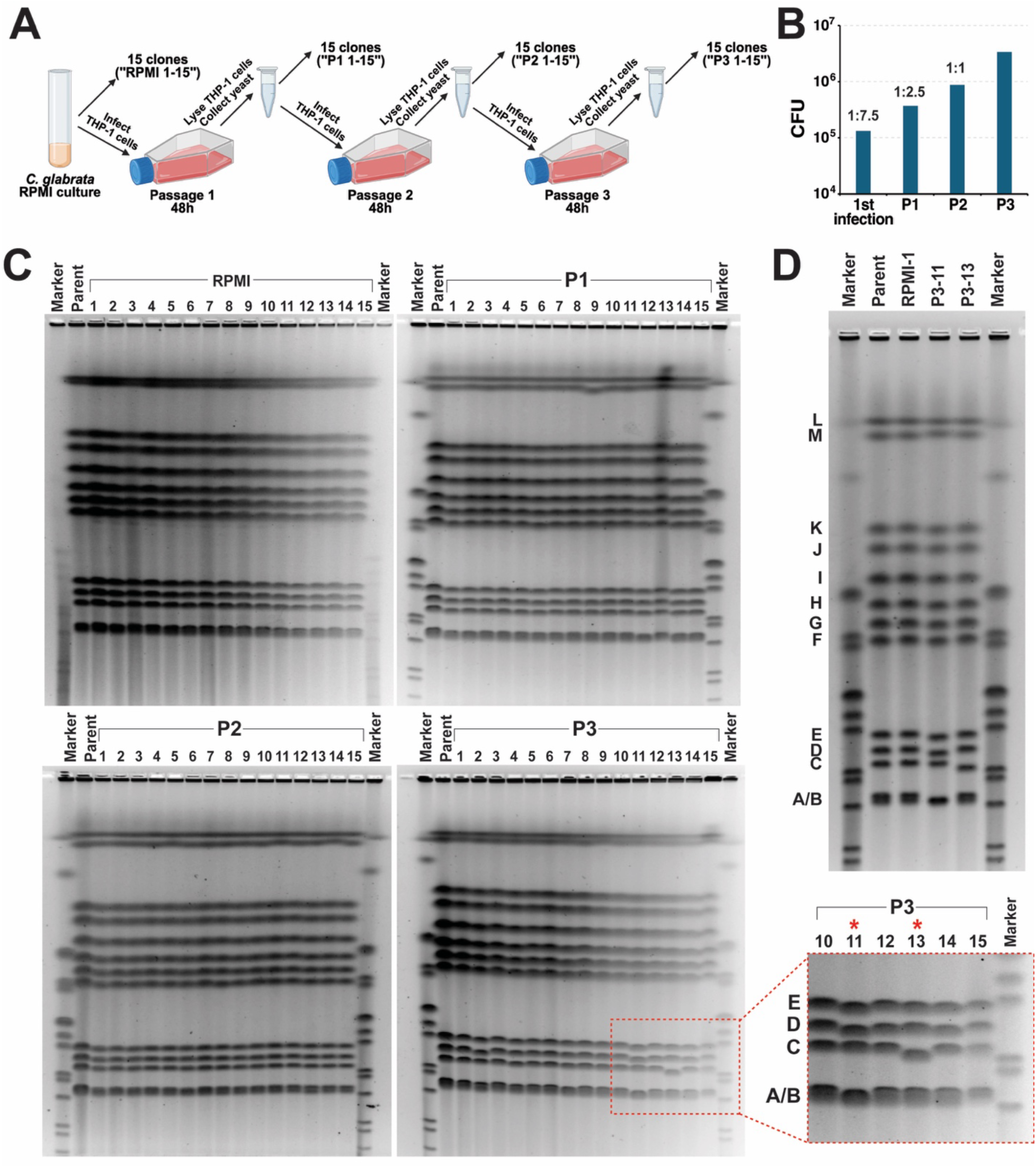
*C. glabrata* cells repeatedly exposed to the intra-macrophage environment have high frequencies of chromosome rearrangements. **A.** A schematic depiction of the macrophage passaging experiment. This panel was created with the help of Biorender (institutional license). **B.** The bar graph shows the CFU used to infect each batch of macrophages (Y-axis), with the corresponding *C. glabrata* : macrophage infection ratio shown above. **C.** PFGE images of 15 clones saved from the initial RPMI culture and 15 clones saved from every passage. Clones P3-11 and P3-13 showed clear chromosomal mobility changes. **D.** Another PFGE experiment using new chromosomal preparations of the parent (CBS138) and clones RPMI-1, P3-11, and P3-13. The same chromosomal alterations were consistently seen in P3-11 and P3-13.

To ask whether intra-macrophage passaging affected chromosome structure, we analyzed all 60 isolates by pulse-field gel electrophoresis (PFGE). Interestingly, whereas none of the isolates from RPMI, passage 1, or passage 2 had obvious chromosomal alterations, two isolates from passage 3 showed clear shortening of chromosomes A/B, D, and E for isolate P3-11 and chromosome C for isolate P3-13 (Fig. 5C). These changes were observed repeatedly using different DNA preparations and different gel runs (e.g., see Fig. 5D). To try to identify the chromosomal regions involved in these alterations, we used Nanopore sequencing of the two strains, as well as of the parental strain CBS138 and strain RPMI-1 as normal controls. Notably, the DNA samples submitted for Nanopore sequencing were the same as those used for PFGE in Fig. 5D. To identify potential chromosomal changes, we used three different methods: a short-read-based approach, a long-read-based approach, and a mixed-model approach that combines both short and long reads for increased accuracy. Surprisingly, none of these analyses identified any detectable chromosomal changes in P3-11 or P3-13 relative to the parental strain or the RPMI isolate.

*C. glabrata* has been noted for carrying an unusually high number of adhesin-like genes, which are thought to be important for virulence, are rich in repeated sequences, and are organized in long tandem repeats, mostly (though not exclusively) found near telomeres (33, 34). These regions are considered to be prone to rearrangements due to non-allelic recombination and have been difficult to resolve, even using long-read sequencing. In fact, the internal (non-telomeric) adhesin gene array on chromosome C remains unresolved in the reference strain CBS138 (35). To determine whether we can obtain other examples of chromosomal structural variants (SV) detectable by PFGE but not by long-read sequencing, we used a set of *C. glabrata* strains whose genomes have been fully assembled by Marcet-Houben et al. (34) using a combination of long-read and short-read technologies. Some of the strains used in that study had been shared by our lab, so they were available to us for PFGE analysis (Fig. S4). A comparison between PFGE results and genome assemblies (34) revealed similarities (Fig. S4 blue chromosome labels) but also clear differences (Fig. S4 red chromosome labels) between the chromosome sizes expected from genome assembly and chromosome mobility. We also consulted supplementary data from Marcet-Houben et al. (34) listing all detected SV in the strains and concluded, again, that it did not explain all observed chromosomal mobility differences.

From these results, we conclude that the altered chromosomal mobility in strains P3-11 and P3-13 is likely due to alterations in the highly repeated regions, which are unresolvable even using state of the art long-read sequencing methods. We also infer that such chromosomal rearrangements occur frequently in intra-macrophage *C. glabrata*, as two out of 15 randomly chosen P3 isolates had distinct altered karyotypes. Finally, it is likely that we are significantly underestimating the extent of SV in macrophage-passaged *C. glabrata*, as PFGE of whole chromosomes can clearly detect only SV that alter chromosome size (such as segmental deletions and duplications, but not inversions) and involve shorter chromosomes (because the mobility of the longer chromosomes would only be visibly affected by very large differences in chromosome size).

### Evidence for evolution of intra-macrophage fitness in *C. glabrata* during macrophage passaging

To ask whether intra-macrophage passaging is also associated with increased sequence diversity, we used Illumina short-read sequencing on all 60 strains from the experiment above, as well our version of the parental strain CBS138. Interestingly, most strains had not acquired additional SNPs relative to the parent, and strains from passage 3 did not have increased SNP frequencies relative to previous passages or to RPMI-derived isolates (Fig. 6A), indicating that intra-macrophage genome instability primarily involves structural rearrangements, most likely at highly repeated sequences. We did identify three different mutations in coding sequences in macrophage-passaged isolates: a missense mutation in *SRC1* in isolates P1-5 and P1-8, a missense mutation in *PRE10* in isolate P2-6, and a frameshift mutation in *RME1* in isolate P2-14 (Fig. 6A). In *S. cerevisiae*, *SRC1* encodes a perinuclear chromosome tethering protein involved in mitotic sister chromatid segregation and rDNA maintenance (36, 37), *PRE10* encodes a subunit of proteasome core complex involved in ubiquitin-dependent and ubiquitin-independent protein degradation (38), and *RME1* encodes a transcriptional regulator of meiosis, mitosis and invasive growth (39, 40).

**Figure 6.**
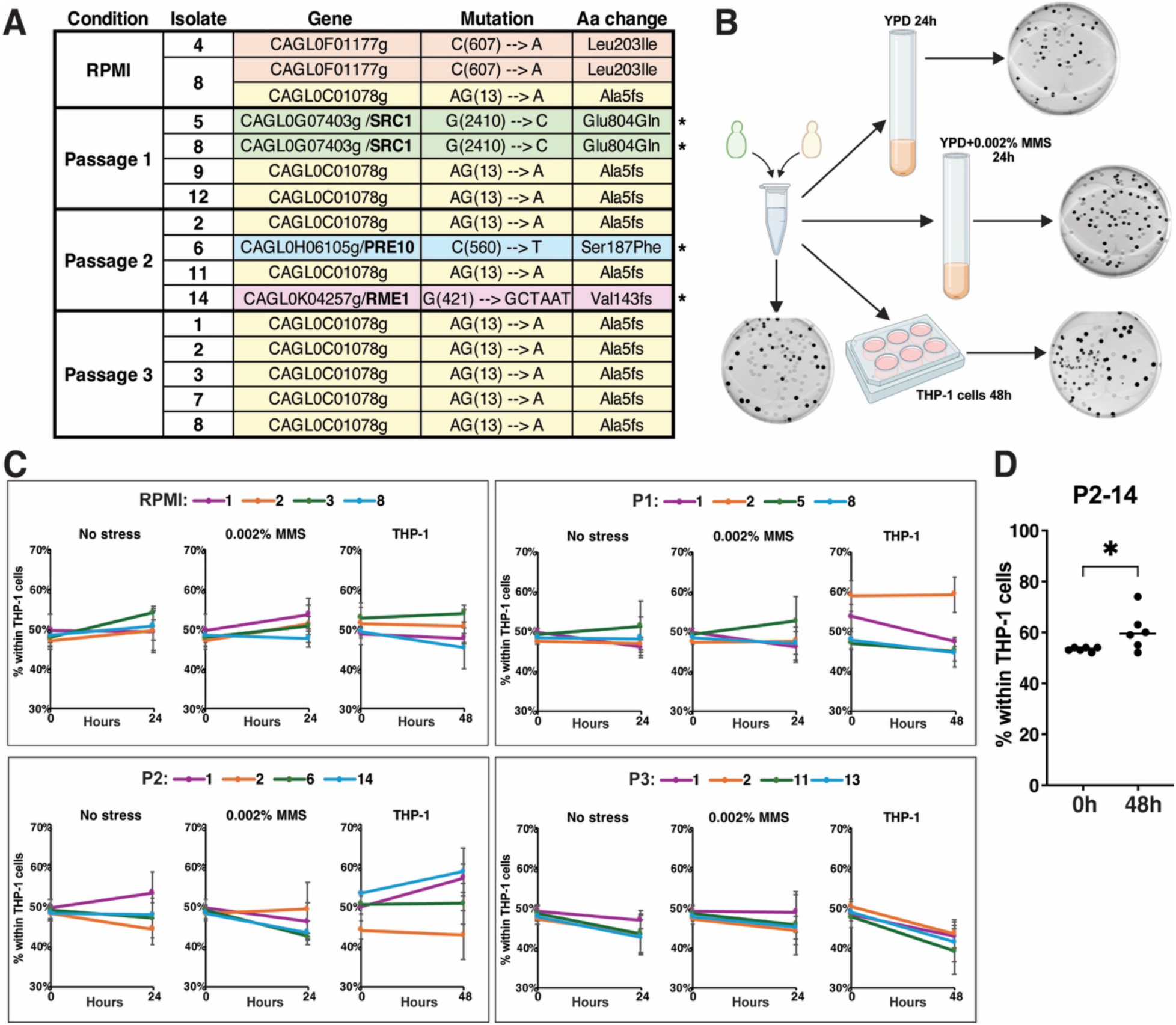
A mutation in a negative regulator of meiosis *RME1* that emerged in *C. glabrata* during macrophage passaging increases *C. glabrata* intra-macrophage competitive fitness. **A.** Summary of coding SNPs identify in macrophage-passaged strains using Illumina short-read sequencing. Asterisks indicate mutations that were confirmed by Sanger sequencing. **B.** A schematic outline of the competition experiments between macrophage-passaged strains and GFP-expressing parental strain CBS138. This panel was created with the help of Biorender (institutional license). **C.** Summary of all competition experiments done with the selected clones from RPMI and passages 1-3. **D.** Clone P2-14 containing the *rme1-Val143fs* mutation outcompetes the parental strain inside macrophages. Each dot represents a biological replicate. The data was normally distributed and the t-test was used for pairwise comparison. * p<0.05

To ask whether the genetic changes (SNPs or large chromosomal changes) occurring in intra-macrophage *C. glabrata* improve its intra-macrophage fitness, we performed a series of competition experiments between selected isolates and the parental strain CBS138 carrying a chromosomally-integrated, constitutively expressed GFP (26). For the competitions, we selected four isolates from each passage, focusing on those carrying either SNPs or chromosomal alterations detectable by PFGE. The competition experiments were performed in culture under two conditions – no stress and low level DNA damage (0.002% MMS) – and in macrophages (Fig. 6B). In each experiment, the two competing strains were mixed at a ratio of 1:1 and either incubated in culture for 24 hours or used to infect macrophages for 48 hours. The relative proportions of the two strains at the beginning and end of each experiment were assessed using plating and imaging of colonies to identify the percentages of fluorescent vs. non-fluorescent colonies (Fig. 6B). These experiments led to the following conclusions. First, successive passages in macrophages did not correlate with progressively improved intra-macrophage fitness (Fig. 6C). In fact, all tested passage 3 isolates showed decreased fitness relative to the parental strain under all three conditions tested. Second, fitness in 0.002% MMS in culture did not correlate with intra-macrophage fitness, consistent with the conclusion that DNA damage was one of several stresses present inside the phagosome. Finally, we found that one passage 2 isolate, P2-14, showed improved fitness inside macrophages but not in culture (Fig. 6C). This was intriguing, so we repeated the macrophage competition experiment with that isolate and obtained a similar result: after 48 hours strain P2-14 consistently comprised a higher fraction of the intra-macrophage population than at the start of the experiment (Fig. 6D). Interestingly, strain P2-14 carried a frameshift mutation in *RME1*, which encodes a zinc finger transcription factor that negatively regulates meiotic gene expression but positively regulates mitotic cell cycle genes and invasive growth (39, 40), raising many new questions about the role of the meiotic apparatus in intracellular *C. glabrata*.

## Discussion

In this study we compared *C. glabrata* transcriptional responses to a diverse panel of *in vitro* stresses and to the intra-macrophage environment. This approach allowed us to identify core groups of *C. glabrata* genes stereotypically induced or repressed by stress and also revealed that although the intra-macrophage environment is not closely recapitulated by any individual *in vitro* stress, among the tested stresses it is most closely resembles amino acid deprivation and DNA damage. The conclusion that *C. glabrata* experiences amino acid deprivation inside macrophages was supported by the phenotype of deleting the master regulator of amino acid biosynthesis *GCN4*, which significantly reduced *C. glabrata* ability to grow in the absence of exogenously supplied amino acids and within macrophages. Similarly, the conclusion that *C. glabrata* experiences DNA damage inside macrophages was supported both by comet assays indicating increased DNA breaks and the fitness defects exhibited by mutants lacking the capacity to repair DNA DSBs. Also consistent with a DNA damaging environment, *C. glabrata* repeatedly passaged through macrophages exhibited a high frequency of chromosomal structural variants (SVs) detected by PFGE. Interestingly, these alterations could not be solved by short- or long-read WGS, strongly suggesting that they involved highly repeated sequences. Finally, one of the SNPs emerging during macrophage-passaging, a frameshift in *RME1* (repressor of <u>me</u>iosis in *S. cerevisiae*), increased *C. glabrata* intra-macrophage fitness. Together, these results shed new light on the environmental milieu and adaptation mechanisms of *C. glabrata* within macrophages, its important host niche, and provide direct evidence for DNA damage in an intra-macrophage pathogen.

Nutrient restriction by phagocytes is a well-appreciated facet of innate immunity, and many studies have shown that the ability of pathogens to compete for nutrient uptake and to synthesize or scavenge nutrients is essential for survival within the host (41, 42). In particular, this was demonstrated for pathogenic fungi and amino acids. For instance, upregulation of amino acid biosynthesis genes has been reported in *C. albicans* taken up by neutrophils (43) and in *C. glabrata* taken up by macrophages by two hours post-infection (7), the latter finding being consistent with our results. Gcn4p is a transcription factor with a conserved role as master regulator of amino acid biosynthesis in response to amino acid deprivation in multiple fungi, including *S. cerevisiae* and *C. albicans* (28, 44). The phenotype of our *C. glabrata gcn4Δ* mutant, which was deficient both in growth without exogenous amino acids and inside macrophages, is consistent with the conclusion that *C. glabrata* experiences amino acid deprivation inside macrophages. However, our data also indicate that the loss of *GCN4* slightly reduces *C. glabrata* survival in the absence of glucose, which may also contribute to its reduced fitness inside macrophages, previously shown to be a glucose-depleted environment (8, 45). Thus, although the most straightforward explanation for our results is that the intra-macrophage environment is depleted of amino acids, other factors may also contribute to the observed phenotypes.

Although it has long been presumed that the intra-macrophage environment damages pathogens’ DNA due to ROS accumulation, direct evidence for this has, to our knowledge, been lacking. Here we optimized the comet assay, a gold-standard tool for DNA break detection, in *C. glabrata* to show that intra-macrophage yeast cells have elevated levels of DNA breaks relative to cultured cells. We used the alkaline comet assay, which detects both single-strand and double-strand DNA breaks (SSB and DSB), with the majority likely being SSB. However, the presence of DSB, which are highly detrimental to genome stability and can be lethal if unrepaired (31, 32), in intra-macrophage *C. glabrata* is suggested by the fitness defects of mutants lacking both HR and NHEJ pathways, which are unable to repair DNA DSB (46). Nevertheless, our data suggest that the DNA lesions occurring in intra-macrophage *C. glabrata* are distinct from those caused by MMS or PHL in culture. First, we show that the latter are largely repaired by HR and that removing NHEJ does not impair cells further; this suggests that MMS- or PHL-induced lesions, but not intra-macrophage lesions, are predominantly repaired using a homologous template, such as the sister chromatid, and may therefore be largely replication-associated (e.g., involving collapsed replication forks). Second, the correlation between transcriptional responses to MMS and to macrophages, while stronger than those for most other examined stresses, was moderate, also suggesting that the nature of the underlying DNA lesions is different. Therefore, the exact structure and context of DNA lesions occurring in intra-macrophage *C. glabrata* is still an open question.

An intriguing aspect of *C. glabrata* biology is the conservation of genes involved in mating and meiosis – two processes never observed in this organism, even though genomic evidence has provided evidence of recombination suggestive of sexual reproduction (47–49). Previously, we showed that a high dose of MMS induces many *C. glabrata* genes whose orthologs are involved in mating and meiosis in *S. cerevisiae* (50). Here, we also show that mating pheromones and pheromone receptors are upregulated as part of the core stress response in *C. glabrata* and that the intra-macrophage environment induces genes whose *S. cerevisiae* orthologs create and repair meiotic DSBs (e.g., *SPO11*, *SPO13*, *SPO22*, *REC8*, and others). Finally, we found that a frameshift mutation in the ortholog of *S. cerevisiae RME1* emerged in *C. glabrata* during macrophage passaging and resulted in improved intra-macrophage fitness. Although *RME1* was originally identified due to its role as a transcriptional repressor of meiotic genes, later it was also shown to promote invasive growth in *S. cerevisiae* (40). Interestingly, in *C. albicans RME1* was shown to promote asexual sporulation driven by nutrient starvation (51), consistent with other examples of transcriptional circuitry rewiring between *S. cerevisiae* and *C. albicans* (52). Although *S. cerevisiae* is much more closely related to *C. glabrata* than to *C. albicans*, transcriptional reprogramming may still have occurred between *S. cerevisiae* and *C. glabrata*, such that *CgRME1* no longer regulates meiosis but other processes. Overall, our results indicate that the genes involved in mating and meiosis in *S. cerevisiae* are upregulated by stress in *C. glabrata*, including within macrophages, but whether this upregulation is relevant to sexual reproduction or is involved in other cellular functions remains to be elucidated.

Karyotype variability is a hallmark feature of clinical *C. glabrata* isolates (17–19), but the circumstances triggering chromosome instability have remained unknown. Previously it was demonstrated that when cells are cultured in laboratory medium (YPD) for up to 70 generations, with the exception of rare supernumerary mini-chromosomes present in some clinical strains but absent in CBS138, all standard *C. glabrata* chromosomes (A-M) remain stable (17). Here we detected visible mobility alterations in chromosomes A/B, C, D, or E in two out of 15 clones obtained from thrice-passaged yeast, and, given that PFGE can more easily identify size variation in smaller chromosomes, the real frequency of structural variation may be even higher. However, because no GCR reporter is available in *C. glabrata*, the precise changes in GCR rate cannot currently be calculated. Interestingly, our previous study demonstrated that the DNA damage checkpoint is significantly attenuated in *C. glabrata* compared to its close relative *S. cerevisiae* (23). Notably, in *S. cerevisiae* mutation of DNA damage checkpoint genes results in high rates of gross chromosomal rearrangements (GCR) (20, 22) but relatively low rates of forward point mutations (53, 54), which is consistent with the effects of macrophage passaging on *C. glabrata*. Given that macrophage infection, proliferation, escape, and re-infection likely occur in host-colonizing *C. glabrata*, these observations likely have clinical relevance.

Our long-read sequencing approach could not identify the structural rearrangements causing the variant karyotypes, suggesting that these alterations involve highly repetitive regions, which in *C. glabrata* are found on all chromosomes and contain copies of various adhesin-like genes thought to be important for virulence (33, 34). Indeed, chromosomal rearrangements due to recombination between repetitive sequences are also a reported consequence of DNA damage and checkpoint defects (55, 56). The effects of adhesin gene rearrangements in *C. glabrata* are not well understood, though it is noteworthy that four out of four thrice-passaged *C. glabrata* strains showed reduced competitive fitness both in macrophages and in rich medium, whereas one twice-passaged strain (the *rme1* mutant) showed improved fitness specifically in macrophages. Intriguingly, our previous study showed that intra-macrophage *C. glabrata* exposed to micafungin develops echinocandin-resistant *fks2* mutations more readily than cultured *C. glabrata* (11). Altogether, our results suggest that the combination of macrophages’ DNA-damaging environment and *C. glabrata*’s attenuated DNA damage checkpoint result in high frequencies of chromosomal alterations mainly involving repeat-rich adhesin genes, with largely detrimental consequences on *C. glabrata* fitness but the occasional emergence of mutants better adapted to their environment, with important implications for the evolution of clinically relevant phenotypes, such as resistance to antifungal drugs or host immunity.

## Materials and Methods

### *C. glabrata* strains, media, and genetic manipulations

Strain CBS138 (a.k.a. ATCC2001) was used for all experiments and to construct all deletion mutants. All strains are listed in Table S2. For genetic manipulations, cells were cultured in standard yeast extract-peptone-dextrose (YPD) medium at 37°C. Deletion mutants were generated in-house using a CRISPR-CAS9 targeted integration replacing the desired ORF by a nourseothricin (NAT)- or hygromycin (HYG)-resistance cassette. The deletion constructs containing the NAT-resistance or HYG-resistance cassette flanked by regions homologous to the locus of interest was either amplified from genomic DNA using ultramer primers (Table S3) or synthesized *de novo* (Twist Bioscience). Integration of the deletion cassettes was performed using CRISPR as described previously (57). Transformants were selected on nourseothricin (NAT)- or hygromycin (HYG)-containing plates and validated by PCR amplification and sequencing of the targeted locus using external primers (Table S3). To create *rad52Δ lig4Δ* double mutants, marker recycling was employed as follows. A shuttle vector (pMVK3) was created by Gibson Assembly (New England Biolabs, USA) combining a *C. glabrata* CEN/ARS backbone from pYC55 (58), a *MET3* promoter sequence from *C. glabrata* CBS138 strain, parts from pSFS3b plasmid (59) containing ORF of ScFLP recombinase, CaACT1 terminator, AgTEF1 promoter and AgTEF1 terminator, and HPH ORF from pFA6a-6xGLY-3xFLAG-hphMX4 (60). *RAD52* was first deleted using a FRT-NAT-FRT cassette with gene-specific flanking arms, synthesized by Twist Bioscience, and confirmed by PCR using primer pairs outside of integration cassette and within the *RAD52* ORF. To recycle the NAT resistance gene, the mutants were transformed with pMVK3 plasmid using hygromycin as a selection marker (800µg/ml in YPD). To induce the flippase, cells were cultured in methionine-deficient synthetic media overnight and plated on YPD plates. Once visible, the colonies were replica-plated on NAT (200 µg/ml) and hygromycin (800 µg/ml) plates. Colonies that were unable to grow both on NAT (marker excision) and hygromycin (plasmid expulsion) were subcloned and FRT-NAT deletion was verified by colony PCR (Emerald AMP MAX, TaKaRa Bio, USA). *LIG4* gene was then removed in the unmarked *rad52Δ* strain using FRT-NAT-FRT cassette using the same approach as above. At least two independent transformants were generated and analyzed for every deletion mutant. Primers were ordered from Integrated DNA Technologies (Coralville, IA, United States) and Azenta (South Plainfield, NJ, United States), and all Sanger sequencing of the above-described constructs was done by Azenta (South Plainfield, NJ, United States). For growth curves, optical density (OD_600_) in the indicated medium was measured hourly using the Tecan Spark Multimode Microplate Reader.

### Exposing *C. glabrata* to stress in culture and harvesting for transcriptomic analysis

For the stress experiments, *C. glabrata* was cultured at 37°C in synthetic medium without amino acids (0.17% yeast nitrogen base without ammonium sulfate or amino acids, 0.5% ammonium sulfate, 2% glucose), with the exception of the amino acid starvation condition, where cells were first cultured in synthetic medium with amino acids (Table S4) and shifted into synthetic medium without amino acids. Overnight cultures were diluted to OD_600_=0.5 and grown for 2 hours at 37°C with shaking, at which point the 0h time-point was collected. For every stress condition, cells were centrifuged, washed with sterile water, resuspended in medium containing the stressor agent, and incubated at 37°C with shaking for the indicated amounts of time. RNA was isolated using the QIAGEN RNEasy Mini Kit (Qiagen cat.74106) according to manufacturer instructions.

### Library Preparation with PolyA selection and Illumina Sequencing

RNA library preparations and sequencing reactions were conducted at Azenta Life Sciences (South Plainfield, NJ, USA) as follows: RNA samples were quantified using Qubit 2.0 Fluorometer (Life Technologies, Carlsbad, CA, USA) and RNA integrity was checked using Agilent TapeStation 4200 (Agilent Technologies, Palo Alto, CA, USA). The RNA sequencing libraries were prepared using the NEBNext Ultra II RNA Library Prep Kit for Illumina using manufacturer’s instructions (New England Biolabs, Ipswich, MA, USA). Briefly, mRNAs were initially enriched with Oligod(T) beads. Enriched mRNAs were fragmented for 15 minutes at 94°C. First strand and second strand cDNA were subsequently synthesized. cDNA fragments were end repaired and adenylated at 3’ends, and universal adapters were ligated to cDNA fragments, followed by index addition and library enrichment by PCR with limited cycles. The sequencing libraries were validated on the Agilent TapeStation (Agilent Technologies, Palo Alto, CA, USA), and quantified by using Qubit 2.0 Fluorometer (ThermoFisher Scientific, Waltham, MA, USA) as well as by quantitative PCR (KAPA Biosystems, Wilmington, MA, USA). The sequencing libraries were clustered on four flowcell lanes. After clustering, the flowcell was loaded on the Illumina HiSeq instrument (4000 or equivalent) according to manufacturer’s instructions. The samples were sequenced using a 2×150bp Paired End (PE) configuration. Image analysis and base calling were conducted by the Control software. Raw sequence data (.bcl files) generated from the sequencer were converted into fastq files and de-multiplexed using Illumina’s bcl2fastq 2.17 software. One mismatch was allowed for index sequence identification. The sequencing data are available from the National Institutes of Health sequence archive under BioProject ID PRJNA1311114.

### RNAseq differential gene expression analysis

FASTQ reads were aligned to *C. glabrata* reference genome CBS138 from the *Candida* genome resource using STAR v *2.7.11b* and quantified using GeneCounts mode. Differential expression analysis was performed using DESeq2 v 3.22. The design formula, expressed as ∼ time + condition + time:condition, allowed for the simultaneous assessment of three effects: the overall effect of time, the effect of the condition at time zero, and the interactive effect of time and condition. Raw counts were used as input, with DESeq2 internally handling normalization via the “median of ratios” method to account for library size and composition biases. Log2 fold-change (LFC) values and adjusted p-values were then extracted for specific contrasts to quantify and assess the significance of the treatment’s effect at different time points relative to the controls.

### *C. glabrata* macrophage infections and serial passaging

Human acute monocyte leukemia cell line-derived macrophages (THP-1; ATCC catalog number TIB-202; Manassas, VA) was used to assess the phagocytosis survival of our *C. glabrata* isolates. RPMI 1640 (Gibco, Fisher Scientific, USA) supplemented with 1% penicillin-streptomycin (Gibco, Fisher Scientific, USA) and 10% heat-inactivated HFBS (Gibco, Fisher Scientific, USA) was used to grow the THP1 cells. One million THP-1 cells treated with 100 nM phorbol 12-myrisate 13-acetate (PMA, Sigma) were seeded into 24-well plates and incubated at 37^°^C in 5% CO_2_ for 48 hours to induce attachment and differentiation into active macrophages. On the day of infection macrophages were washed with PBS, fresh RPMI was added, and *C. glabrata* cells were added to macrophages. Subsequently, the plates were centrifuged (200xg, 1 minute) and incubated at 37^°^C in 5% CO_2_. After 3 hours, the supernatant RPMI containing unengulfed cells was removed, the pellets were washed five times with phosphate buffered saline (PBS), and fresh RPMI was added. To extract intra-macrophage *C. glabrata* at 3/24/48 hour time-points, macrophages were lysed by adding 1 ml ice-cold water and shaking for 45 min. The yeast obtained from lysed macrophages was diluted and plated on YPD for CFU enumeration or processed for additional experiments, such as comet assays. *C. glabrata* intra-macrophage fitness was calculated as the fold-increase in intra-macrophage *C. glabrata* between 3h and 24h post-infection, normalized to the wild-type control average from the same experiment. For serial macrophage passaging, *C. glabrata* cells were cultured in RPMI, harvested, and used to infect THP-1 cells as described above. Dilutions of the RPMI culture were plated on YPD for CFU counts, and 15 random colonies from the YPD plates were saved (isolates RPMI1-15). 48 hours after infection, the macrophages were lysed, the *C. glabrata* cells collected and then immediately used to re-infect a fresh batch of THP-1 macrophages. At the same time, dilutions were plated on YPD to obtain CFU counts, and 15 randomly chosen colonies were saved (isolates P1-1 through P1-15 from passage 1). This was repeated twice more to obtain 15 isolates from passage 2 (P2-1 through P2-15) and 15 isolates from passage 3 (P3-1 through P3-15).

### Comet assay

*C. glabrata* cells were resuspended in sterile Buffer S (1M Sorbitol, 25mM KH_2_PO_4_, pH 6.5) containing 1.5% SeaPlaque low-melting agarose (Lonza), 1 mg/ml Zymolyase 100T (US Biological), and 50uM β-mercaptoethanol (Fisher Chemical). 10 µl of this cell suspension (maintained at 37°C to keep from solidifying) was then pipetted onto glass slides precoated with 0.5% regular agarose (Promega), immediately covered with a coverslip, and placed in a humidity chamber at 37°C for 30 mins to allow cell wall digestion. The slides were then moved to 4°C for 15 minutes to allow the agarose to solidify, the coverslips were carefully removed, and the slides were immersed in ice-cold lysis solution (30mM NaOH, 1M NaCl, 0.05% (w/v) lauroysarcosine, 50mM EDTA, 10mM Tris-HCl, pH 10) for 20 mins at 4°C. Slides were then placed in the electrophoresis slide tray containing ice-cold Alkaline Electrophoresis Solution (8 g of NaOH pellets and 2 mL of 500mM EDTA pH 8 in 1 liter of MilliQ water, pH>13) and voltage was applied for 10 min at 10V in CometAssay Electrophoresis System II (Bio-techne, #4250-050-ES). Next, the slides were washed twice in sterile water and once in 70% ethanol, 5 minutes each. The slides were dried at 30°C for 20 minutes (or until fully dry). Comets were stained for 30 minutes using a solution of propidium iodide (2 µg/ml) in TBE buffer, rinsed several times with water, covered with a coverslip, sealed, and imaged on a Nikon Eclipse Ti2 inverted microscope with a Hamamatsu ORCA-Flash4.0 camera using the TRITC filter at 60X magnification. OpenComet (30) was used to analyze the comet images using background correction and the “brightest region” setting recommended for cells with moderate DNA damage.

### Pulse-field gel electrophoresis

*C. glabrata* full-length chromosomal DNA was prepared and separated using a procedure modified from *S. cerevisiae* conditions described earlier (61). 5 ml *C. glabrata* cultures were grown overnight at 37°C, and 1 ml aliquots of such saturated cultures were centrifuged at 3,000 rpm for 5 minutes. The supernatant was carefully removed by aspiration, yielding fresh cell pellets of ∼30 mg (wet weight). Pellets were gently resuspended in 810 μl of molten (45C) low melting point agarose solution (0.5% in 100 mM EDTA pH 7.5) with additional 36 μl of Zymolyase 20T 25 mg/ml solution and immediately transferred to BioRad PFGE plug molds. Agarose-embedded samples were incubated overnight at 37°C for cell wall digestion, then treated with sarcosyl and proteinase K solution at 50°C for 5 hours. Plugs were cleaned through successive washes in TE buffer and then resolved in a BioRad CHEF DRII instrument with the following running buffer and settings: 0.5x TBE buffer at 14°C, 180 ml 1% PFGE-grade agarose gel in portrait orientation (21 cm long by 14 cm wide), 5 Volts/cm, initial and final switch times of 65 and 175 seconds with linear ramping, respectively, and total run time of 66 hours. Gels were stained with ethidium bromide, destained in water, then imaged with a BioRad UV transillumator. Additional details of the optimized *C. glabrata* PFGE protocol are available upon request.

### Whole genome sequencing (WGS) of macrophage-passaged *C. glabrata*

Genomic DNA from *C. glabrata* was sequenced using short-read Illumina (for parent strain CBS138 as well as all RPMI and macrophage passaged clones) and long-read Nanopore (for parent strain, one selected RPMI clone, and two selected macrophage-passaged clones) WGS as described previously (62, 63). YeaStar Genomic DNA kit was used to extract high quality genomic DNA samples, which were then processed for short-read Illumina multiplex WGS libraries using seqWell plexWell 96. DNA for long-read Nanopore WGS was prepared using as starting material plugs prepared for PFGE (see above; chromosome-length genomic DNA embedded in agarose). DNA was extracted from PFGE plugs using the procedure established by Oxford Nanopore (https://nanoporetech.com/document/extraction-method/agarose-plug-dna). The sequencing data are available from the National Institutes of Health sequence archive under BioProject ID PRJNA1311114.

### Identification of SNPs in macrophage-passaged *C. glabrata*

We implemented a bioinformatics pipeline in python 3.11, which combined STAR (v2.7.6a) for read alignment and SAMtools (v1.16.1) and BCFtools (v1.16.1) for variant calling. Illumina reads were aligned to the *C. glabrata* CBS138 reference genome using STAR, and resulting BAM files were then sorted and indexed using samtools sort and samtools index respectively. A pileup file was generated from the sorted BAM files using samtools mpileup -A -d 500. Finally, BCFtools was used to call variants from the pileup data with the call -m -v command, producing a VCF file containing all identified SNPs and small indels.

### Identification of structural variation in macrophage-passaged *C. glabrata*

To identify structural variants, we used short-read, long-read, and combination methods. For the short-read method we used Pythor, a Python-based tool, to identify changes in copy number variation (CNV). The process involved aligning the short sequencing reads to the reference genome using STAR. The aligned BAM files where then processed by Pythor to generate read depth profiles across the genome. By comparing these profiles to a control sample, Pythor could identify regions with significant increases or decreases in read depth, which correspond to gene duplications or deletions, respectively. This approach leverages the high accuracy and throughput of Illumina sequencing to provide a precise, genome-wide view of CNVs. For the long-read analysis, we used Nanopore reads to perform de novo assembly and identify large-scale structural variants. This method involved two main steps: first, the long reads were assembled into a contiguous genome using Flye and Canu. Then, the quality of the assembly was assessed using QUAST (Quality Assessment Tool for Genome Assemblies). QUAST generates a comprehensive report on assembly metrics, including contig length, genome size, and the number of genes, allowing us to identify large-scale chromosomal rearrangements such as translocations or inversions that would be difficult to detect with Illumina reads. Finally we developed a mixed-model pipeline that integrated both short and long reads to enhance the detection of structural variants. This pipeline involved several steps: 1) Alignment: Long read alignment waas done using minimap2 and combined with the existing short-read BAM files generated by the STAR alignment. 2) Calling: We then used two different structural variant (SV) callers to leverage the unique strengths of each read type. SVIM (Structural Variant Identification using Minimap2) was used to call SVs from the long-read alignments and Manta was used to call SVs from the short-read alignments. 3) Combine: Output VCF files from SVIM and Manta were then merged using SURVIVOR. This step combined the predictions from both callers, reducing false positives and creating a more comprehensive list of high-confidence Svs. 4) Manual Validation: The merged list of variants was validated using read depth analysis of the short-read data. We checked the coverage at the sites of predicted deletions and were able to confirm that a drop in read depth was associated with the deletion.

### Statistical analysis

All data were analyzed using GraphPad Prism 10 software. Pairwise comparisons were performed using an unpaired t-test or the Mann-Whitney test, depending on the normality of distribution. Normality was determined using the Shapiro-Wilk test. P values of ≤0.05 were considered statistically significant.

## Acknowledgements

This work was supported by 5R01AI109025 to DSP and ES, 1R21AI168729 to ES, and 5R35GM119788 to JLA. MH was supported as fellow through the Predoctoral Training in Quantitative Cell & Molecular Biology award 5T32GM132057. We thank Dr. Jane Usher for valuable advice regarding the comet assay methodology.

---

**Figure S1.**
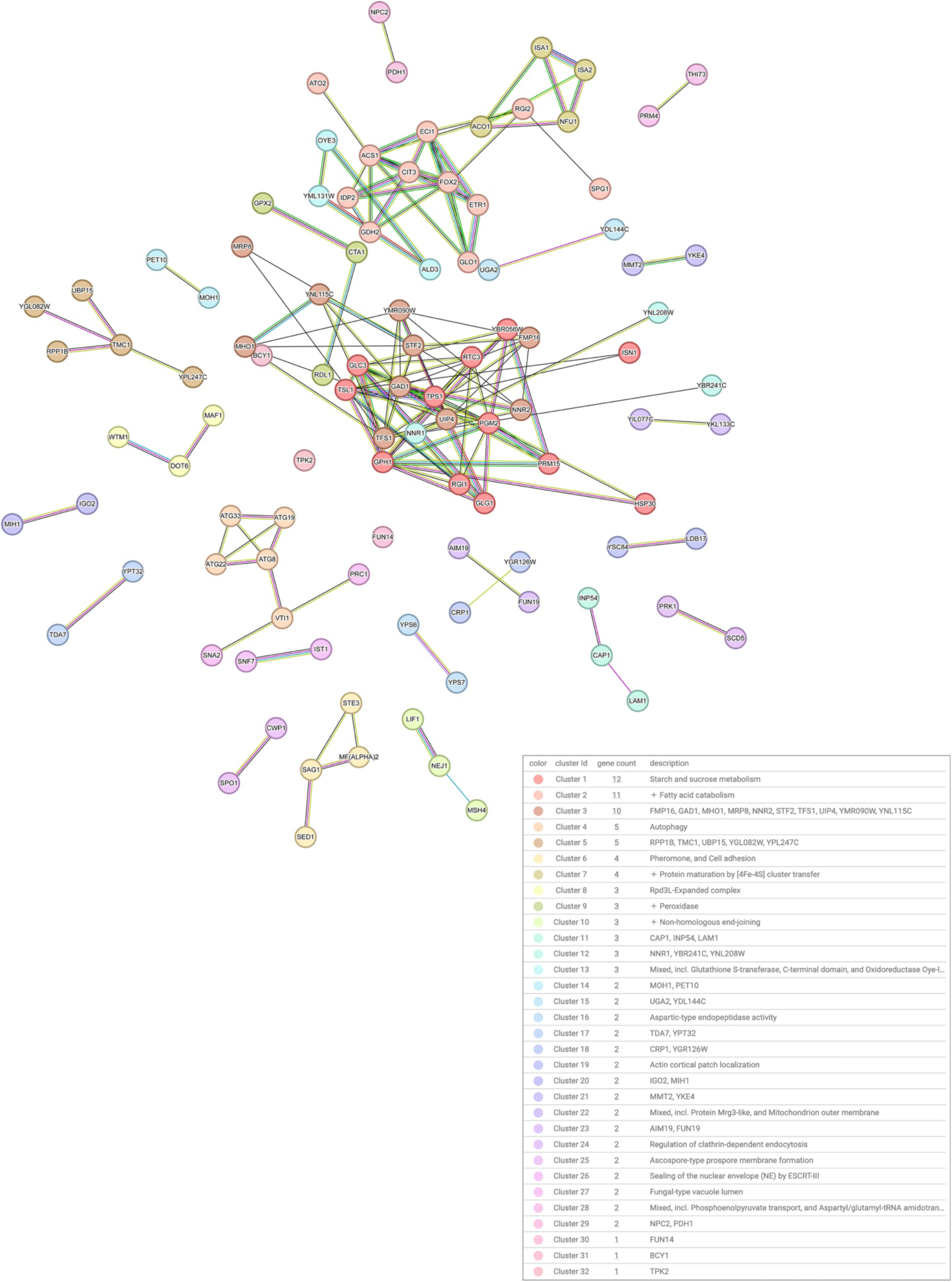
String-DB network of the core set of genes upregulated by all examined environmental stresses. The String-DB clustering algorithm was used with medium confidence (0.4) setting, omitting unconnected nodes, and using the default MCL clustering inflation parameter 3.

**Figure S2.**
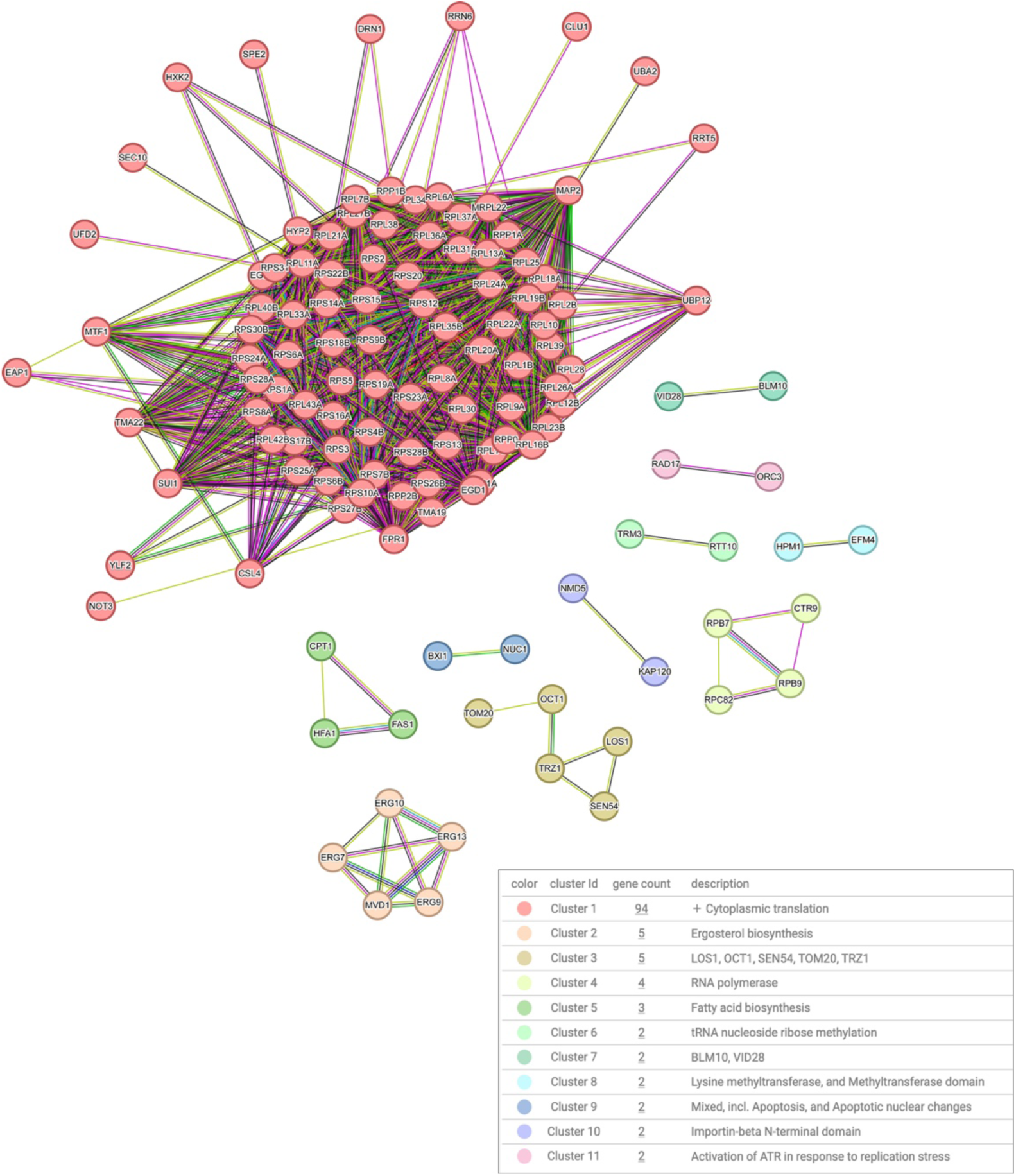
String-DB network of the core set of genes downregulated by all examined environmental stresses. The String-DB clustering algorithm was used with medium confidence (0.4) setting, omitting unconnected nodes, and using the default MCL clustering inflation parameter 3.

**Figure S3.**
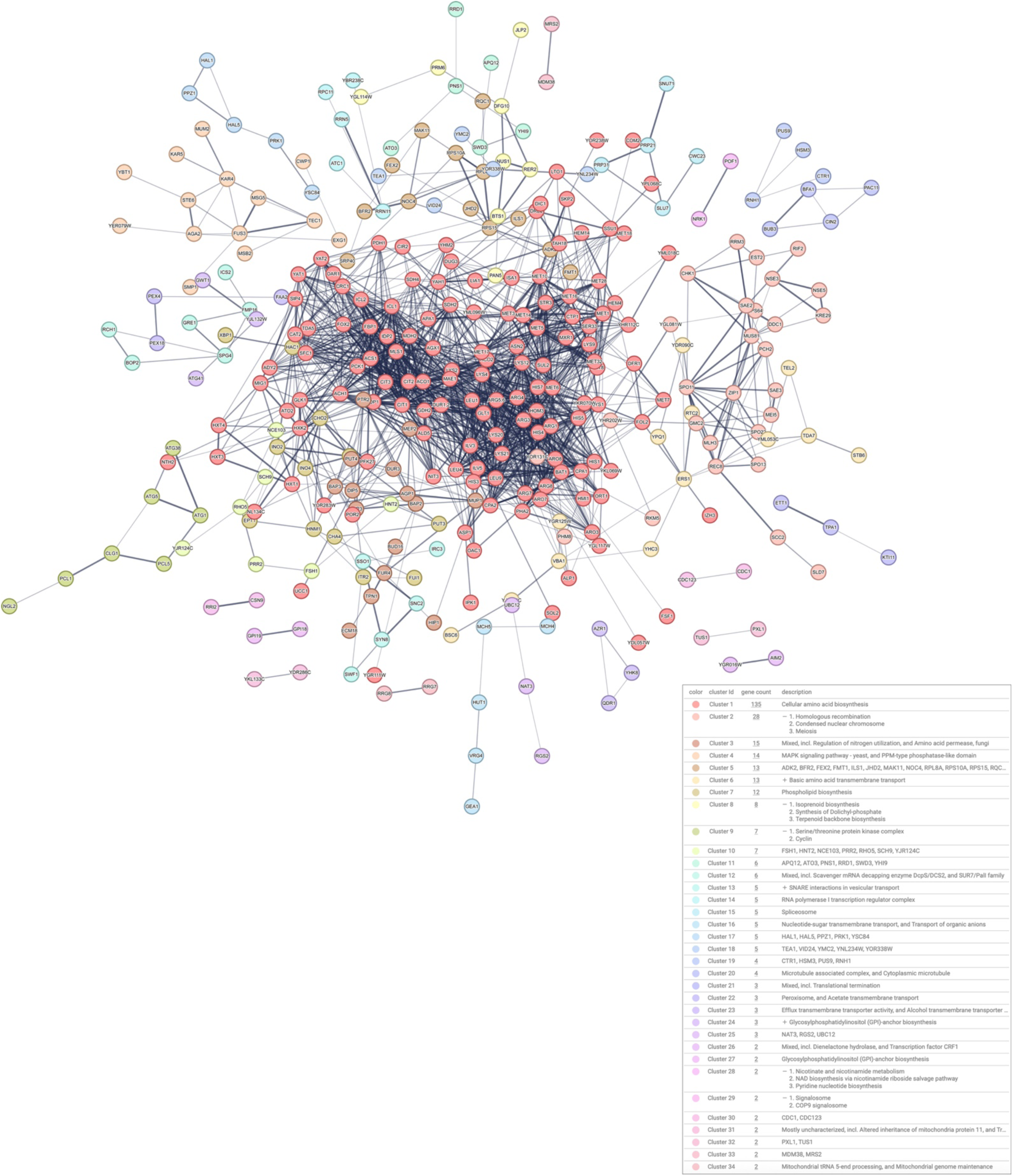
String-DB network of the core set of genes upregulated (log2FC>1, FDR p<0.05) at 3 hours post-macrophage infection. The String-DB clustering algorithm was used with medium confidence (0.4) setting, omitting unconnected nodes, and using MCL clustering inflation parameter 1.5 to reduce the number of detected clusters (because of the large number of genes).

**Figure S4.**
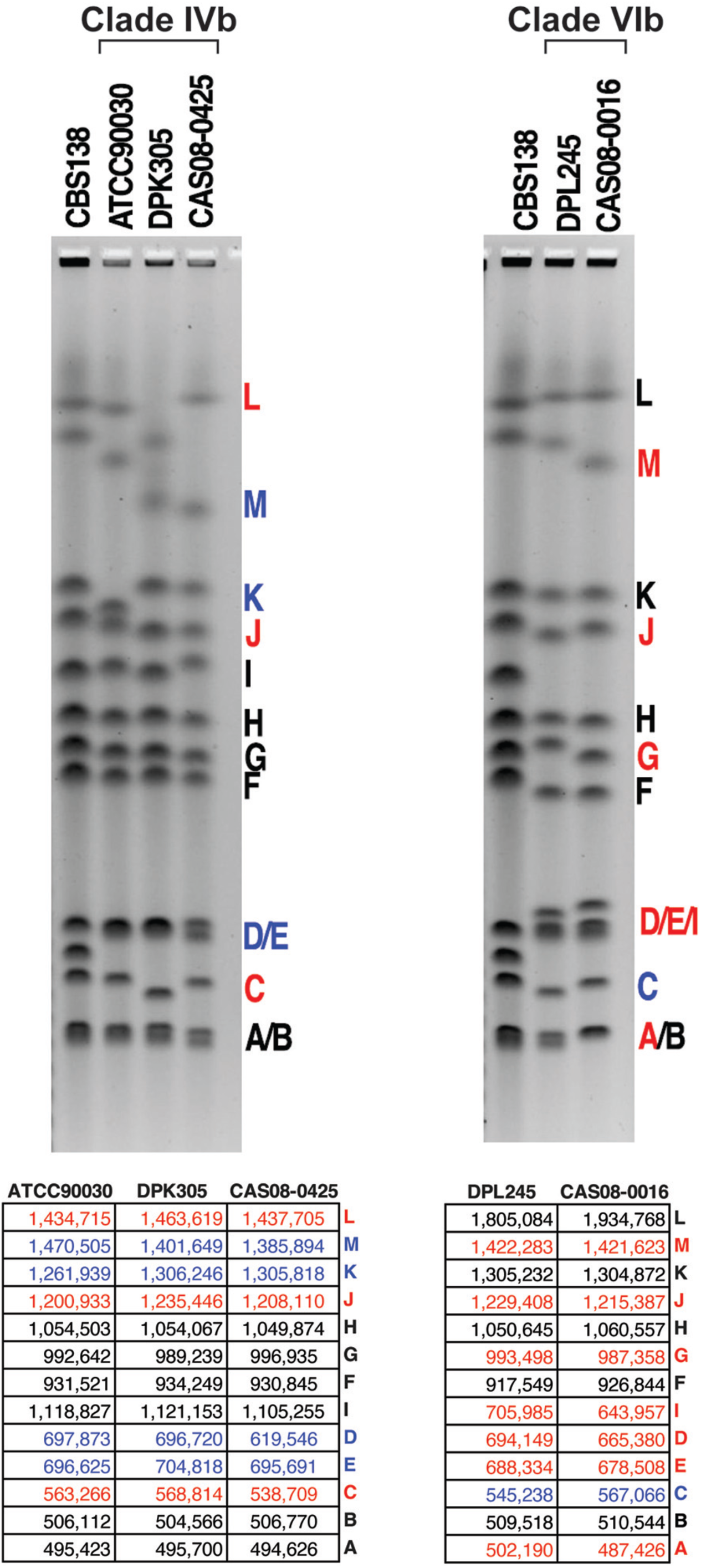
PFGE analysis of strains whose genomes were assembled using a combination of long- and short-read sequences. In Marcet-Houben et al. (34). The tables show the expected chromosome sizes based on the genome assemblies in (34). Similarities between PFGE mobility and expected size are shown in blue, whereas differences are shown in red.

## REFERENCES

1. Denning DW. Global incidence and mortality of severe fungal disease. Lancet Infect Dis. 2024.

2. Sonnberger J, Kasper L, Lange T, Brunke S, Hube B. “We’ve got to get out“-Strategies of human pathogenic fungi to escape from phagocytes. Mol Microbiol. 2024;121(3):341–58.

3. Erwig LP, Gow NA. Interactions of fungal pathogens with phagocytes. Nat Rev Microbiol. 2016;14(3):163–76.

4. Lorenz MC, Bender JA, Fink GR. Transcriptional response of Candida albicans upon internalization by macrophages. Eukaryot Cell. 2004;3(5):1076–87.

5. Miramón P, Pountain AW, Lorenz MC. *Candida auris*-macrophage cellular interactions and transcriptional response. Infect Immun. 2023:e0027423.

6. Kaur R, Ma B, Cormack BP. A family of glycosylphosphatidylinositol-linked aspartyl proteases is required for virulence of Candida glabrata. Proc Natl Acad Sci U S A. 2007;104(18):7628–33.

7. Rai MN, Lan Q, Parsania C, Rai R, Shirgaonkar N, Chen R, et al. Temporal transcriptional response of C*andida glabrata* during macrophage infection reveals a multifaceted transcriptional regulator CgXbp1 important for macrophage response and fluconazole resistance. Elife. 2024;13.

8. Shen Q, Ray SC, Evans HM, Deepe GS, Rappleye CA. Metabolism of Gluconeogenic Substrates by an Intracellular Fungal Pathogen Circumvents Nutritional Limitations within Macrophages. mBio. 2020;11(2).

9. Healey KR, Perlin DS. Fungal Resistance to Echinocandins and the MDR Phenomenon in Candida glabrata. J Fungi (Basel). 2018;4(3).

10. Perlin DS, Rautemaa-Richardson R, Alastruey-Izquierdo A. The global problem of antifungal resistance: prevalence, mechanisms, and management. Lancet Infect Dis. 2017;17(12):e383–e92.

11. Arastehfar A, Daneshnia F, Cabrera N, Penalva-Lopez S, Sarathy J, Zimmerman M, et al. Macrophage internalization creates a multidrug-tolerant fungal persister reservoir and facilitates the emergence of drug resistance. Nat Commun. 2023;14(1):1183.

12. Kasper L, Seider K, Hube B. Intracellular survival of Candida glabrata in macrophages: immune evasion and persistence. FEMS Yeast Res. 2015;15(5):fov042.

13. Rai MN, Balusu S, Gorityala N, Dandu L, Kaur R. Functional genomic analysis of Candida glabrata-macrophage interaction: role of chromatin remodeling in virulence. PLoS Pathog. 2012;8(8):e1002863.

14. Forche A, Cromie G, Gerstein AC, Solis NV, Pisithkul T, Srifa W, et al. Rapid Phenotypic and Genotypic Diversification After Exposure to the Oral Host Niche in *Candida albicans*. Genetics. 2018;209(3):725–41.

15. Ene IV, Farrer RA, Hirakawa MP, Agwamba K, Cuomo CA, Bennett RJ. Global analysis of mutations driving microevolution of a heterozygous diploid fungal pathogen. Proc Natl Acad Sci U S A. 2018;115(37):E8688–E97.

16. Ford CB, Funt JM, Abbey D, Issi L, Guiducci C, Martinez DA, et al. The evolution of drug resistance in clinical isolates of Candida albicans. Elife. 2015;4:e00662.

17. Polakova S, Blume C, Zarate JA, Mentel M, Jorck-Ramberg D, Stenderup J, et al. Formation of new chromosomes as a virulence mechanism in yeast Candida glabrata. Proc Natl Acad Sci U S A. 2009;106(8):2688–93.

18. Healey KR, Jimenez Ortigosa C, Shor E, Perlin DS. Genetic Drivers of Multidrug Resistance in Candida glabrata. Front Microbiol. 2016;7:1995.

19. Shin JH, Chae MJ, Song JW, Jung SI, Cho D, Kee SJ, et al. Changes in karyotype and azole susceptibility of sequential bloodstream isolates from patients with Candida glabrata candidemia. J Clin Microbiol. 2007;45(8):2385–91.

20. Myung K, Chen C, Kolodner RD. Multiple pathways cooperate in the suppression of genome instability in Saccharomyces cerevisiae. Nature. 2001;411(6841):1073–6.

21. Putnam CD, Srivatsan A, Nene RV, Martinez SL, Clotfelter SP, Bell SN, et al. A genetic network that suppresses genome rearrangements in Saccharomyces cerevisiae and contains defects in cancers. Nat Commun. 2016;7:11256.

22. Myung K, Datta A, Kolodner RD. Suppression of spontaneous chromosomal rearrangements by S phase checkpoint functions in Saccharomyces cerevisiae. Cell. 2001;104(3):397–408.

23. Shor E, Garcia-Rubio R, DeGregorio L, Perlin DS. A Noncanonical DNA Damage Checkpoint Response in a Major Fungal Pathogen. mBio. 2020;11(6).

24. Gasch AP, Spellman PT, Kao CM, Carmel-Harel O, Eisen MB, Storz G, et al. Genomic expression programs in the response of yeast cells to environmental changes. Mol Biol Cell. 2000;11(12):4241–57.

25. Cuellar-Cruz M, Briones-Martin-del-Campo M, Canas-Villamar I, Montalvo-Arredondo J, Riego-Ruiz L, Castano I, et al. High resistance to oxidative stress in the fungal pathogen Candida glabrata is mediated by a single catalase, Cta1p, and is controlled by the transcription factors Yap1p, Skn7p, Msn2p, and Msn4p. Eukaryot Cell. 2008;7(5):814–25.

26. Arastehfar A, Daneshnia F, Hovhannisyan H, Fuentes D, Cabrera N, Quinteros C, et al. Overlooked *Candida glabrata* petites are echinocandin tolerant, induce host inflammatory responses, and display poor *in vivo* fitness. mBio. 2023;14(5):e0118023.

27. Arastehfar A, Daneshnia F, Hovhannisyan H, Cabrera N, Ilkit M, Desai JV, et al. A multidimensional assessment of in-host fitness costs of drug resistance in the opportunistic fungal pathogen Candida glabrata. FEMS Yeast Res. 2024;24.

28. Hinnebusch AG. Translational regulation of GCN4 and the general amino acid control of yeast. Annu Rev Microbiol. 2005;59:407–50.

29. Collins A, Møller P, Gajski G, Vodenková S, Abdulwahed A, Anderson D, et al. Measuring DNA modifications with the comet assay: a compendium of protocols. Nat Protoc. 2023;18(3):929–89.

30. Gyori BM, Venkatachalam G, Thiagarajan PS, Hsu D, Clement MV. OpenComet: an automated tool for comet assay image analysis. Redox Biol. 2014;2:457–65.

31. Cannan WJ, Pederson DS. Mechanisms and Consequences of Double-Strand DNA Break Formation in Chromatin. J Cell Physiol. 2016;231(1):3–14.

32. Waterman DP, Haber JE, Smolka MB. Checkpoint Responses to DNA Double-Strand Breaks. Annu Rev Biochem. 2020;89:103–33.

33. Xu Z, Green B, Benoit N, Sobel JD, Schatz MC, Wheelan S, et al. Cell wall protein variation, break-induced replication, and subtelomere dynamics in Candida glabrata. Mol Microbiol. 2021;116(1):260–76.

34. Marcet-Houben M, Alvarado M, Ksiezopolska E, Saus E, de Groot PWJ, Gabaldón T. Chromosome-level assemblies from diverse clades reveal limited structural and gene content variation in the genome of Candida glabrata. BMC Biol. 2022;20(1):226.

35. Xu Z, Green B, Benoit N, Schatz M, Wheelan S, Cormack B. De novo genome assembly of Candida glabrata reveals cell wall protein complement and structure of dispersed tandem repeat arrays. Mol Microbiol. 2020;113(6):1209–24.

36. Rodríguez-Navarro S, Igual JC, Pérez-Ortín JE. SRC1: an intron-containing yeast gene involved in sister chromatid segregation. Yeast. 2002;19(1):43–54.

37. Mekhail K, Seebacher J, Gygi SP, Moazed D. Role for perinuclear chromosome tethering in maintenance of genome stability. Nature. 2008;456(7222):667–70.

38. Geng F, Tansey WP. Similar temporal and spatial recruitment of native 19S and 20S proteasome subunits to transcriptionally active chromatin. Proc Natl Acad Sci U S A. 2012;109(16):6060–5.

39. Toone WM, Johnson AL, Banks GR, Toyn JH, Stuart D, Wittenberg C, et al. Rme1, a negative regulator of meiosis, is also a positive activator of G1 cyclin gene expression. EMBO J. 1995;14(23):5824–32.

40. van Dyk D, Hansson G, Pretorius IS, Bauer FF. Cellular diperentiation in response to nutrient availability: The repressor of meiosis, Rme1p, positively regulates invasive growth in Saccharomyces cerevisiae. Genetics. 2003;165(3):1045–58.

41. Zhang YJ, Rubin EJ. Feast or famine: the host-pathogen battle over amino acids. Cell Microbiol. 2013;15(7):1079–87.

42. Traven A, Naderer T. Central metabolic interactions of immune cells and microbes: prospects for defeating infections. EMBO Rep. 2019;20(7):e47995.

43. Rubin-Bejerano I, Fraser I, Grisafi P, Fink GR. Phagocytosis by neutrophils induces an amino acid deprivation response in Saccharomyces cerevisiae and Candida albicans. Proc Natl Acad Sci U S A. 2003;100(19):11007–12.

44. Tripathi G, Wiltshire C, Macaskill S, Tournu H, Budge S, Brown AJ. Gcn4 co-ordinates morphogenetic and metabolic responses to amino acid starvation in Candida albicans. EMBO J. 2002;21(20):5448–56.

45. Williams RB, Lorenz MC. Multiple Alternative Carbon Pathways Combine To Promote Candida albicans Stress Resistance, Immune Interactions, and Virulence. mBio. 2020;11(1).

46. Mehta A, Haber JE. Sources of DNA double-strand breaks and models of recombinational DNA repair. Cold Spring Harb Perspect Biol. 2014;6(9):a016428.

47. Carrete L, Ksiezopolska E, Pegueroles C, Gomez-Molero E, Saus E, Iraola-Guzman S, et al. Patterns of Genomic Variation in the Opportunistic Pathogen Candida glabrata Suggest the Existence of Mating and a Secondary Association with Humans. Curr Biol. 2018;28(1):15–27 e7.

48. Muller H, Hennequin C, Gallaud J, Dujon B, Fairhead C. The asexual yeast Candida glabrata maintains distinct a and alpha haploid mating types. Eukaryot Cell. 2008;7(5):848–58.

49. Wong S, Fares MA, Zimmermann W, Butler G, Wolfe KH. Evidence from comparative genomics for a complete sexual cycle in the ‘asexual’ pathogenic yeast Candida glabrata. Genome Biol. 2003;4(2):R10.

50. Shor E, Perlin DS. DNA damage response of major fungal pathogen Candida glabrata opers clues to explain its genetic diversity. Curr Genet. 2021;67(3):439–45.

51. Hernández-Cervantes A, Znaidi S, van Wijlick L, Denega I, Basso V, Ropars J, et al. A conserved regulator controls asexual sporulation in the fungal pathogen Candida albicans. Nat Commun. 2020;11(1):6224.

52. Johnson AD. The rewiring of transcription circuits in evolution. Curr Opin Genet Dev. 2017;47:121–7.

53. Cordón-Preciado V, Ufano S, Bueno A. Limiting amounts of budding yeast Rad53 S-phase checkpoint activity results in increased resistance to DNA alkylation damage. Nucleic Acids Res. 2006;34(20):5852–62.

54. Fasullo M, Tsaponina O, Sun M, Chabes A. Elevated dNTP levels suppress hyper-recombination in Saccharomyces cerevisiae S-phase checkpoint mutants. Nucleic Acids Res. 2010;38(4):1195–203.

55. Mieczkowski PA, Lemoine FJ, Petes TD. Recombination between retrotransposons as a source of chromosome rearrangements in the yeast Saccharomyces cerevisiae. DNA Repair (Amst). 2006;5(9-10):1010–20.

56. George CM, Alani E. Multiple cellular mechanisms prevent chromosomal rearrangements involving repetitive DNA. Crit Rev Biochem Mol Biol. 2012;47(3):297–313.

57. Grahl N, Demers EG, Crocker AW, Hogan DA. Use of RNA-Protein Complexes for Genome Editing in Non-albicans Candida Species. mSphere. 2017;2(3).

58. Yanez-Carrillo P, Orta-Zavalza E, Gutierrez-Escobedo G, Patron-Soberano A, De Las Penas A, Castano I. Expression vectors for C-terminal fusions with fluorescent proteins and epitope tags in Candida glabrata. Fungal Genet Biol. 2015;80:43–52.

59. Tscherner M, Stappler E, Hnisz D, Kuchler K. The histone acetyltransferase Hat1 facilitates DNA damage repair and morphogenesis in Candida albicans. Mol Microbiol. 2012;86(5):1197–214.

60. Funakoshi M, Hochstrasser M. Small epitope-linker modules for PCR-based C-terminal tagging in Saccharomyces cerevisiae. Yeast. 2009;26(3):185–92.

61. Zhang H, Zeidler AF, Song W, Puccia CM, Malc E, Greenwell PW, et al. Gene copy-number variation in haploid and diploid strains of the yeast Saccharomyces cerevisiae. Genetics. 2013;193(3):785–801.

62. Heasley LR, Watson RA, Argueso JL. Punctuated Aneuploidization of the Budding Yeast Genome. Genetics. 2020;216(1):43–50.

63. Heasley LR, Argueso JL. Genomic characterization of a wild diploid isolate of Saccharomyces cerevisiae reveals an extensive and dynamic landscape of structural variation. Genetics. 2022;220(3).

64. Nolte H, MacVicar TD, Tellkamp F, Krüger M. Instant Clue: A Software Suite for Interactive Data Visualization and Analysis. Sci Rep. 2018;8(1):12648.

65. Ge SX, Jung D, Yao R. ShinyGO: a graphical gene-set enrichment tool for animals and plants. Bioinformatics. 2020;36(8):2628–9.

